# CRISPR-RNA binding drives structural ordering that primes Cas7-11 for target cleavage

**DOI:** 10.1101/2024.08.01.606276

**Authors:** Calvin P. Lin, Harry Li, Daniel J. Brogan, Tianqi Wang, Omar S. Akbari, Elizabeth A. Komives

## Abstract

Type III-E CRISPR-Cas effectors, referred to as Cas7-11 of gRAMPs, are single proteins that cleave target RNAs without nonspecific collateral cleavage, opening new possibilities for RNA editing. Here, biochemical assays combined with amide hydrogen-deuterium exchange (HDX-MS) experiments provide a first glimpse of the conformational dynamics of apo Cas7-11. HDX-MS revealed the backbone comprising the four Cas7 zinc-binding RRM folds are well-folded but insertion sequences are highly dynamic and fold upon binding crRNA. The crRNA causes folding of disordered catalytic loops and β-hairpins, stronger interactions at domain-domain interfaces, and folding of the Cas7.1 processing site. Target RNA binding causes only minor ordering around the catalytic loops of Cas7.2 and Cas7.3. We show that Cas7-11 cannot independently process the CRISPR array and that binding of partially processed crRNA induces multiple states in Cas7-11 and reduces target RNA cleavage. The insertion domain shows the most ordering upon binding of mature crRNA. Finally, we show a crRNA-induced conformational change in one of the TPR-CHAT binding sites providing an explanation for why crRNA binding facilitates TPR-CHAT binding. The results provide the first glimpse of the apo state of Cas7-11 and reveal how its structure and function are regulated by crRNA binding.

## INTRODUCTION

CRISPR Cas systems, composed of various CRISPR Cas effectors, provide adaptive immunity against invading viruses by interfering with invading nucleic acids (1,2). Three types of Cas effectors utilize a programmable CRISPR RNA (crRNA) to recognize and cleave complementary single-stranded target RNA (tgRNA). The multi-protein type III effectors, type III-A (Csm) and type III-B (Cmr), assemble through multiple Cas7 subunits that bind crRNA and cleave ssRNA (3). The Type VI systems, Cas13a-d, are smaller single-protein effectors with bilobed architecture that target and cleave ssRNA which activates additional non-specific collateral RNA cleavage (4–6). While Cas13’s off-target collateral cleavage activity has proven useful in developing RNA-sensing diagnostic tools (7,8), it hinders its application as a gene editing tool due to cell toxicity and organismal lethality (9,10). The recently discovered subtype III-E CRISPR Cas effectors, Cas7-11 or gRAMP (giant <u>R</u>epeat-<u>A</u>ssociated <u>M</u>ysterious <u>P</u>rotein), are RNA targeting single-protein effectors that have potential in RNA manipulation and therapeutic tools (11,12). Cas7-11 allows for specific RNA target cleavage without collateral off-target RNA degradation and minimal cell toxicity relative to Cas13. Recently, effective trans-splicing of endogenous RNA by Cas7-11 in human cells has been demonstrated (13,14).

As a descendant of type III-A/B effectors; the type III-E effector, Cas7-11, share structural and mechanistic elements, particularly within the Cas7 subunits. The evenly spaced Cas7 subunits of type III effectors sport an RNA recognition motif (RRM) βαββαβ fold and a thumb-like β-hairpin that inserts into the crRNA-tgRNA duplex, resulting in flipping of adjacent bases in both strands (15,16). A nearby catalytic loop containing a conserved Asp is positioned 3’ to the flipped target base to hydrolyze the 2’OH group, resulting in a 6-nt target cleavage interval (3,17).

Cas7-11 is composed of Cas7 domains, (Cas7.1-7.4), a Cas11 domain, the insertion domain (INS), and a Carboxy-terminal extension domain (CTE) that are fused together through 4 linkers (**Figure 1A and D**) (18). Cas7.1, processes the pre-crRNA by cleaving it at position-15 (**Figure 1B**) and recognizes and linearizes the 5’-tag of the direct repeat (DR) (**Figure 1C**), while Cas7.1-7.4 and the INS recognize the spacer region of the crRNA and the complementary tgRNA (19–21). The Cas11 domain has been shown to be necessary for positioning the tgRNA for cleavage (18,20,21). Cas7.2 and Cas7.3 have thumb-like β-hairpins responsible for base-flipping in both RNA strands and a conserved Asp residue responsible for tgRNA cleavage, resulting in cleavage in a 6-nt interval 3’ to the flipped 4^th^ and 10^th^ nucleotides (18). The Cas7.4 domain lacks a β-hairpin and an associated base-flip, and the INS domain is inserted within the domain. The insertion (INS) domain binds the top of the RNA duplex, although additional functions of the INS remain unknown (18,21).

**Figure 1.**
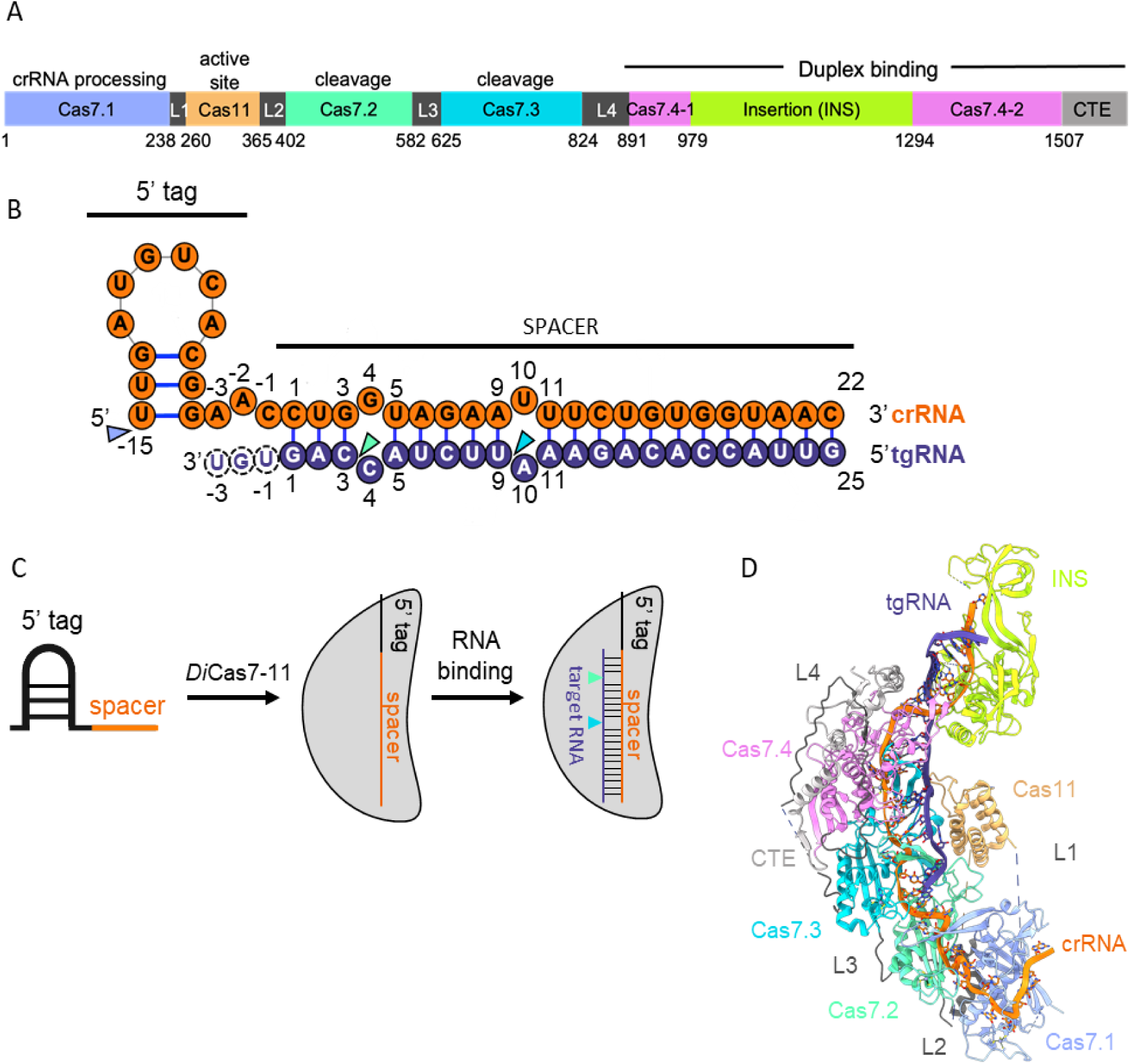
Overview of crRNA, tgRNA, and *Desulfonema ishimotonii* Cas7-11 (*Di*Cas7-11) used in HDX-MS experiments. (**A**) Domain organization of *Di*Cas7-11 labeled by their respective functions. (**B**) Upon *Di*Cas7-11 binding, crRNA is flipped at bases -2, 4, and 10, while tgRNA is flipped at bases 4 and 10. crRNA processing site and tgRNA cleavage sites are denoted by colored by the domain where the respective catalytic residue is found. Target bases -1 to -3, shown with dashed outlines, do not base pair. (**C**) Cartoon diagram of cleavage mechanism. The 5’-tag stem loop loses secondary structure upon association with *Di*Cas7-11. crRNA allows for target recognition through base pairing and subsequent cleavage. (**D**) Structure of *Di*Cas7-11-crRNA-tgRNA (PDB:7WAH) showing the four Cas7 domains (Cas7.1, Cas7.2, Cas7.3, and Cas7.4), the Cas11 domain, the carboxy-terminal extension (CTE) domain, and the INS connected by linkers.

Several mechanistic questions arise from differences between Cas7-11 and other type III effectors. *Desulfonema ishimotonii* Cas7-11 (*Di*Cas7-11) is putatively self-sufficient for crRNA processing (18,19), but *Sb*Cas7-11 does not process its own pre-crRNA (19) and type III-A/B Cas protein complexes process pre-crRNA using ancillary nucleases such as Cas6 or RNase E (22,23). The novel insertion domain has not been assigned a role beyond duplex binding. The insertion domain of the type III-Dv Cas effectors, a partially fused type III evolutionary intermediate between type III-A/B and type III-E Cas effectors, was shown to “seed” tgRNA binding (24). Type III-A/B/E Cas effectors have not shown to have such “seeding” capabilities (3,16), although one *Sb*Cas7-11 study suggests a directionality of tgRNA binding (20).

To gain insight into this new subtype of Cas proteins, we carried out assays to determine whether *Di*Cas7-11 could process CRISPR arrays and which form of the crRNA was best utilized by *Di*Cas7-11 for tgRNA cleavage. We also performed comprehensive HDX-MS experiments to characterize the protein dynamics of *Di*Cas7-11 to characterize the structure of the apo state (for which no structure is available) and to understand how crRNA and tgRNA binding alter the protein dynamics. We report here that the Cas7 RRM folds are well-structured but most of the rest of the protein is only weakly-folded in the absence of crRNA. Binding of crRNA markedly orders the protein backbone in regions directly contacting the crRNA and in parts of the protein that are further from the crRNA binding site such as the Cas11 domain. The dynamics of the INS vary depending on which crRNA is bound and whether tgRNA is bound. Finally, the HDX-MS data reports on regions of *Di*Cas7-11 that could not be resolved in the structure such as those responsible for binding TPR-CHAT.

## MATERIALS AND METHODS

### Protein Expression and Purification

To generate plasmids for recombinant protein expression, ORFs of WT *Di*Cas7-11, d*Di*Cas7-11(D429A/D654A), and *Di*Cas7-11ΔINS(979–1293) from plasmids used for mammalian cell culture were subcloned into a modified pET28a vector containing an N-terminal 6xHis-SUMO tag as previously described (25).

Plasmids were transformed into Rosetta™2(DE3) pLysS Competent Cells (Novagen). Cells were grown in 1L LB supplemented with 50 mg/L kanamycin and 34 mg/L chloramphenicol at 37°C until OD_600_ reached ∼1.0. After incubation on ice for 15 minutes, expression was induced by addition of 0.1 mM IPTG followed by continued growth at 18°C for 20 h. Cells were pelleted and resuspended in a lysis buffer containing 50 mM Tris-HCl pH 8.0, 300 mM NaCl, 10 mM imidazole, 5% glycerol (v/v), and 3 mM β-mercaptoethanol supplemented with Protease Inhibitor Cocktail (Sigma P2714), 5 mM PMSF, and 2.5 U/mL salt active nuclease (Sigma SRE0015). Cells were lysed via sonication and clarified by centrifugation at 25000 xg for 30 minutes. His-tagged protein was bound to 3 mL HisPur™ Ni-NTA Resin (Thermo 88221) equilibrated with lysis buffer in a glass Econo-Column® (Bio-Rad) by flowing through the clarified lysate. The column was washed with 12 column volume (CV) of wash buffer (50 mM Tris-HCl pH 8.0, 500 mM NaCl, 30 mM imidazole, and 5% glycerol) followed by elution with 7 CV of elution buffer (50 mM Tris-HCl pH 8.0, 300 mM NaCl, 300 mM imidazole, and 5% glycerol). The 6xHis-SUMO tag was removed by dialyzing the eluate overnight at 4°C with ∼0.6 mg of 6xHis-tagged Ulp1 (Yeast SUMO Protease) (26) against a dialysis buffer (20 mM HEPES-NaOH pH 7.5, 250 mM NaCl, 5% glycerol, and 1 mM DTT). Ulp1, the cleaved tag, and additional impurities were removed by flowing the dialyzed sample onto the same Ni-NTA column equilibrated in 20mM HEPES-NaOH pH 7.5, 250 mM NaCl, 25 mM imidazole, 5% glycerol, and 1 mM DTT. Cation exchange chromatography was performed on 2x1-mL HiTrap Heparin HP Column (Cytiva) with a NaCl gradient from 200-1000 mM NaCl followed by gel filtration chromatography on a HiLoad 16/600 Superdex 200 column equilibrated in 20 mM HEPES-NaOH pH 7.5, 600 mM NaCl, 5% glycerol (v/v), 2 mM DTT on an ÄKTA Pure (Cytiva) at 4°C. Fractions were analyzed by SDS-PAGE and pure fractions were selected to be concentrated to ∼10 µM and stored in small aliquots at -80°C for future use.

### Assays

#### In vitro cleavage assays

*In vitro* crRNA processing and tgRNA cleavage assays were performed in 10 µL reactions with the following final concentrations: 40 U Murine RNase Inhibitor (NEB), and 6 mM MgCl_2_. Other reagents were added as needed at the following concentrations; 500 nM *Di*Cas7-11, 125 nM crRNA/pre-crRNA, 240 nM 6-FAM 5’-labeled target RNA (ordered from IDT), and 1 U *E. coli* ribonuclease III (RNase III, Ambion Inc.). The target RNA was ordered from IDT and is reported in **Table S1**. Reactions were supplemented with either 1 µL 10X reaction buffer (200 mM HEPES-NaOH pH 7.5, 600 mM NaCl) or 1 µL of manufacturer-provided 10X RNase III Reaction buffer (100 mM Tris pH 7.9, 500 mM NaCl, 100 mM MgCl_2_, and 10 mM DTT) when using RNase III. crRNA processing reactions were incubated for 60 min at 37°C and target cleavage reactions were incubated for 90 minutes at 37°C. Upon completion, reactions were denatured with 2X RNA dye (NEB B0363S) at 95°C for 10 min. Samples (10 µL) were then analyzed on a pre-run (200 V for 60 min) 15% TBE-Urea PAGE gel (BioRad 4566055) at 200 V for 35 min. Gels were stained with SYBR Gold (Invitrogen S11494) and incubated for 10 min at room temperature on a shaker and washed in 1X TBE for 10 min on a shaker before imaging with an ENDURO^TM^ GDS (Labnet).

#### Electrophoretic Mobility Shift Assays

Ribonucleoprotein complexes (RNPs) were pre-formed by mixing Cas7-11 variants and crRNA at 1:1 final concentration ratio for 20 min before adding to an EMSA master mix (Final EMSA contained 40 mM HEPES pH 7.5, 100 mM NaCl, 5% glycerol, 1 mM DTT, 4 U Murine Rnase Inhibitor (NEB), 5 mM EDTA, 2.5 µg/mL heparin, and 10 nM 40-nt 6-FAM 5’-labeled tgRNA (IDT)). Samples were incubated at 37°C for 20 min followed by 20 min at room temperature. Samples were loaded onto a 10% TBE Gel and run at 4°C for 36 minutes at 180V. Gels were imaged using a G:Box chemi XX6 (SynGene).

*In vitro* transcription for crRNA production

The *Di*Cas7-11 single array crRNA (IDT) is reported in **Table S1**. All other crRNAs were produced from dsDNA templates with a T7 promoter as previously described (24). dsDNA templates were generated from template-less PCR and purified with MinElute PCR Purification Kit (Qiagen 28004). To convert dsDNA to ssRNA crRNA; *in vitro* transcription was performed using the MEGAScript^TM^ T7 Transcription Kit (Invitrogen AM1334). ssRNAs were then purified using the MEGAClear^TM^ Transcription Clean-Up Kit (Invitrogen AM1908).

### Hydrogen-Deuterium Exchange Mass Spectrometry

*Di*Cas7-11 or the inactive mutant d*Di*Cas7-11 (D429A/D654A) were prepared as described above with cation exchange followed by gel filtration in H_2_O-based HDX buffer composed of 20 mM HEPES pH 7.5, 150 mM NaCl, 5% glycerol, 1 mM DTT. crRNA_37_, crRNA_57_, and 25-nt target RNA (**Table S1**, IDT) were resuspended in HDX buffer. To form the crRNA-protein complex, protein was incubated with a 2.5 molar excess of crRNA for 1.5 h at 4°C. To form the tgRNA-crRNA-protein complex, 3.75 molar excess tgRNA (to protein), was added to the preformed crRNA-protein complex and incubated for an additional 1 h at 4°C. To prepare the D_2_O buffer, the H_2_O HDX buffer was subjected to speed vac evaporation and resuspended an equal volume of D_2_O.

HDX-MS experiments were performed using a Waters Synapt G2-Si time-of-flight mass spectrometer equipped with a nanoACQUITY UPLC system with HDX technology and a LEAP PAL liquid handling system. Samples (4 µL) were incubated for 5 min at 25°C and then mixed with 56 µL H_2_O buffer, as a control; or D_2_O buffer for 30 s, 60 s, or 120 s. Controls and time points were performed in triplicate. Deuterium exchange was quenched by mixing 50 µL of the protein sample with 60 µL of quench buffer (3 M Guanidine HCl, 2.2% formic acid) and incubated for 2 min at 1°C (the pH of the mixed protein-quench solution was measured to be 2.70 at 0°C). The quenched solution (50 µL) was injected onto an in-line pepsin column (immobilized pepsin, Pierce Inc.) at 15°C and resulting peptides were captured on a BEH C18 Vanguard precolumn before separation on a C18 column (Acquity UPLC BEH C18, 1.7 µM, 1.0 x 50 mm, Waters Corporation) using a 7 to 85% acetonitrile gradient in 0.1% formic over 7.5 min and directly electrosprayed into the mass spectrometer. The mass spectrometer was set to collect data in the Mobility ESI+ mode, with mass acquisition range of 200 to 2000 (m/z) and scan time 0.4 s. Continuous lock mass correction was accomplished with infusions of leu-enkephalin (m/z = 556.2771) every 30 s (mass accuracy of 1 ppm for calibration standard). For peptide identification, the mass spectrometer was set to collect data in Mobility MS^E^, ESI+ mode instead.

Peptides were identified from triplicate MS^E^ analyses of 7 µM *Di*Cas7-11 or d*Di*Cas7-11 samples using PLGS 2.5 (Waters Corporation). Peptide masses were identified using a minimum number of 250 ion counts for low-energy peptides and 50 ion counts for their fragment ions. Peptides identified in PLGS were then analyzed in DynamX 3.0 (Waters Corporation) with initial filters as follows; cutoff score of 6.8, minimum products per amino acid of 0.2, maximum MH+ error tolerance of 5 ppm, and retention time standard deviation of 5%. All mass envelopes were manually checked. The deuterium uptake for each peptide was calculated by comparing the centroids of the mass envelopes of the deuterated samples versus the undeuterated controls following previously established methods and deuterium uptake was corrected for back exchange as previously described (27). Deuterium uptake plots, coverage maps, calculated fractional uptake differences, and PDB structures overlayed with HDX data were generated using DECA - github.com/komiveslab/DECA (28). The uptake plots y-axis maximum is reflective of the total possible amides that can exchange within the peptide and fitted with an exponential curve for ease of viewing using DECA. Community guidelines were followed and are reported in **Table S2**, and the data are publicly available on the Massive data repository (29).

## RESULTS

### HDX-MS reveals that crRNA binding structures Cas7-11

No apo Cas7-11 structures are currently available. To understand the structure and dynamics of the apo Cas7-11, Cas7-11 bound to crRNA, and Cas7-11 bound to crRNA and tgRNA, we performed HDX-MS experiments on the apo WT *Di*Cas7-11 with and without crRNA. Target RNA binding was probed using a catalytically inactive mutant d*Di*Cas7-11-crRNA with and without tgRNA. HDX-MS measures foldedness and solvent accessibility of protein backbones (27,30). Following deuterium incubation, reactions were quenched and proteins were proteolyzed by pepsin for bottom-up protein identification and HDX analysis. Peptic peptides covered 94.4% of the 1601 amino acid wild-type protein and 92.1% of the inactive mutant (**Supplementary Figure S1 and S2**) allowing study of every domain including protein segments not resolved in cryo-EM structures of WT *Di*Cas7-11-crRNA (PDB: 7YN9) (21) and d*Di*Cas7-11-crRNA-tgRNA (PDB:7WAH) (18) (**Supplementary Figure S3**). The structure of *Di*Cas7-11-crRNA with the crRNA hidden was used to represent the apo state as there is no structure of apo *Di*Cas7-11. The apo protein showed a wide range of fractional deuterium uptake (FU) with much of the concave crRNA binding surface being highly exchanging (**Figure 2A**), while the convex side comprised of the RRM folds had low deuterium uptake. Notably, a pattern was observed in which the thumb-like β-hairpins and catalytic loops exhibited high FU, indicating high solvent accessibility. In contrast, the core of the RRM fold (β1-α1-β2-β3-α2-β4), except the RRM fold of Cas7.1, showed low FU in the apo state, suggesting they were well-structured prior to crRNA binding in the apo state (**Figure 2A**) (18). The two active sites, the crRNA processing site, the INS domain, and the Cas11 domain all had highly exchanging regions. To compare HDX between different bound states, fractional uptake differences (FUD) at 60 s deuterium incubation were determined. Upon crRNA addition, marked decreases in exchange (≥ -30% FUD) were observed throughout the protein (**Figure 2B**). Many highly exchanging regions became very low exchanging in the crRNA-bound state. These exchange differences cannot be explained solely by crRNA interaction and likely indicate folding/structuring of *Di*Cas7-11 upon crRNA binding. Subsequent addition of the tgRNA caused few additional decreases in uptake which were localized to the Cas7.2 and Cas7.3 active site, while the INS region showed an increase in deuterium uptake upon tgRNA binding (**Figure 2C**). The Cas7.4 domain showed lower FUD changes compared to other Cas7 domains. The minimal differences in HDX upon tgRNA binding supports previous studies that showed no structural change from target binding (21).

**Figure 2.**
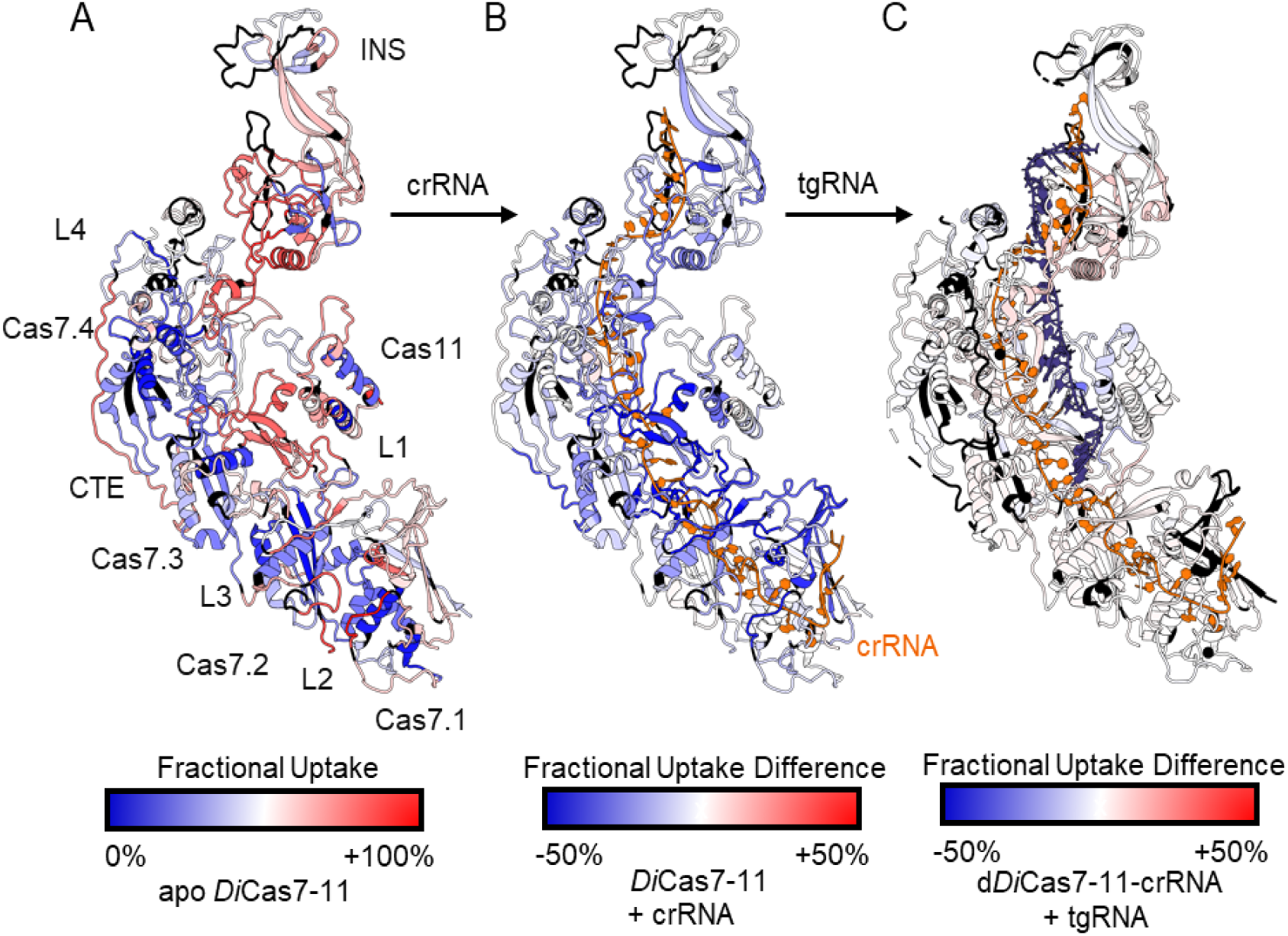
HDX-MS reveals solvent accessibility changes arising from RNA binding. (**A**) Fractional uptake changes were mapped to a structure of WT crRNA-bound *Di*Cas7-11 (PDB:7YN9) with crRNA hidden to represent apo *Di*Cas7-11. The structure is colored according to a scale from blue indicating 0% deuterium uptake to red indicating 100% deuterium uptake. All data are 60 s deuterium incubation and averages of technical triplicates. Regions in black were not covered by peptides from the HDX-MS experiments. (**B**) Fractional uptake differences (FUD), upon crRNA addition to *Di*Cas7-11, were mapped to the same structure as in **A** with crRNA (shown in orange). The structure was colored according to FUD on a scale from blue indicating a 50% reduction in deuterium uptake upon crRNA binding to red indicating a 50% increase in deuterium uptake. (**C**) FUD upon tgRNA (shown in dark slate blue) binding to d*Di*Cas7-11-crRNA mapped onto structure of d*Di*Cas7-11-crRNA-tgRNA (PDB:7WAH).

### crRNA binding induces structuring of the pre-crRNA processing site

Prior to ribonucleoprotein (RNP) complex formation between Cas effectors and their evolved crRNA, crRNAs must be processed following CRISPR array transcription (31). *Di*Cas7-11, is somewhat unique, as it is able to bind and process the 35-nt 5’ direct repeat (DR) of pre-crRNA to a 15-nt 5’-tag and by cleaving off an additional 20-nt stem-loop (11). The Cas7.1 pre-crRNA processing site showed pronounced decreases in deuterium uptake ranging from -30% to - 47.6% FUD (**Figure 3A and B**). Exchange decreases were localized to the region surrounding the scissile phosphate, including the β-strands of the RRM fold. The β1 strand of Cas7.1 represented by the peptide corresponding to residues 39-60, containing the pre-crRNA processing active site residue H43 (18), underwent a marked decrease in deuterium uptake (-40.1% FUD) (**Figure 3B**, black arrow). Neighboring regions of Cas7.1, covered by peptides corresponding to residues 11-17, 33-39, 99-106, 147-170, and 147-173 also showed significant decreases in deuterium uptake (FUDs of -46.0%, -47.6% -42.4%, -31.0%, -58.3% respectively) (**Figure 3B**). These large decreases in exchange cannot be due only to RNA interaction and likely reflect structuring of Cas7.1 upon pre-crRNA binding. By comparing the HDX-MS results for two peptides corresponding to residues 147-170 and 147-173, we deduced that residues 171-173 of the β-hairpin undergo complete protection, likely due to interaction with the flipped base(18,19,21,32).

**Figure 3.**
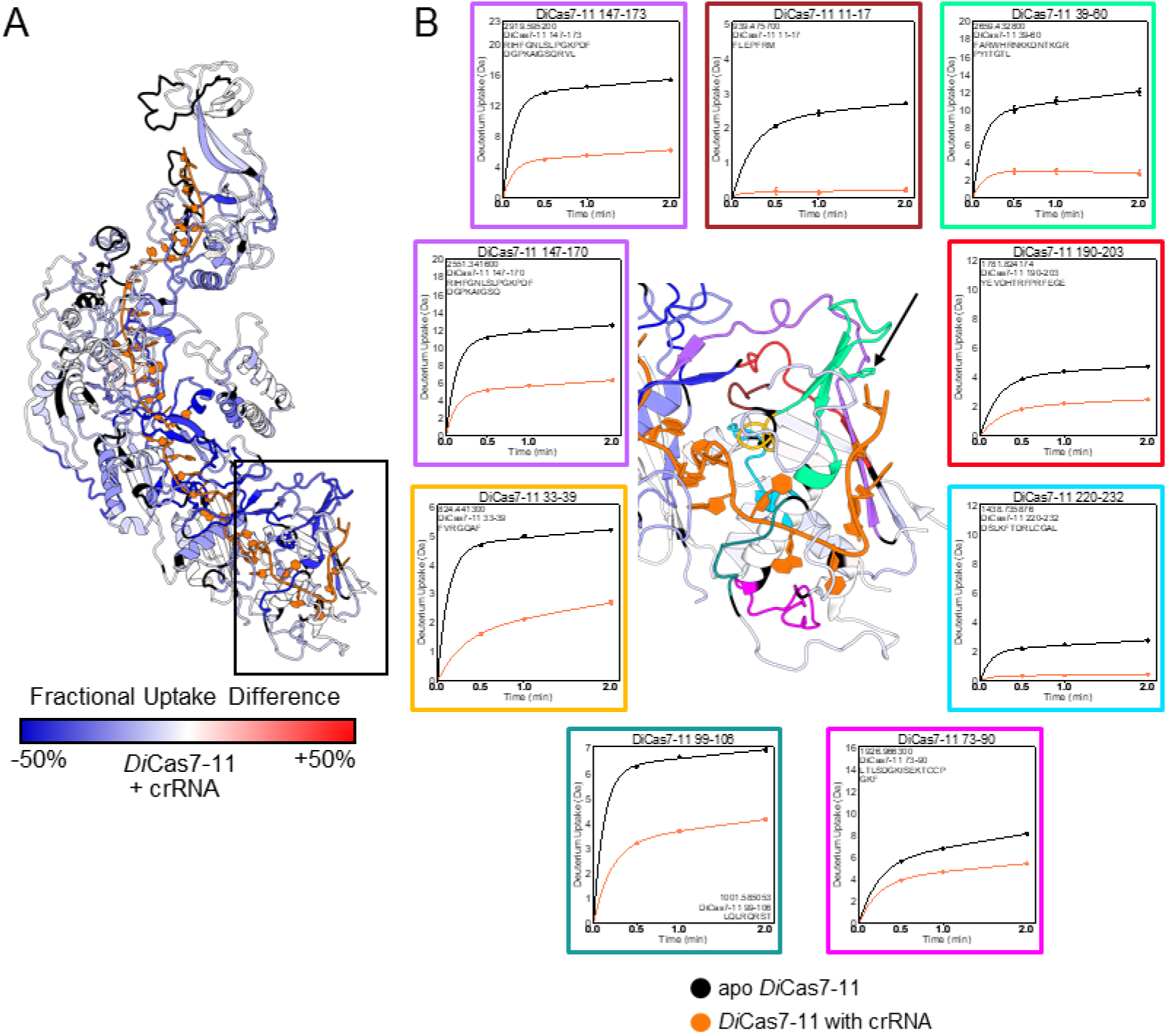
HDX-MS reveals crRNA-binding effects in the processing site of *Di*Cas7-11. (**A**) FUD upon crRNA binding mapped to *Di*Cas7-11-crRNA structure (PDB:7YN9) with inset box around Cas7.1. (**B**) Zoomed-in inset of Cas7.1 domain colored by corresponding HDX uptake plots. Other regions are colored according to FUD, while regions not detected by MS are shown in black. Data is shown as mean values ± SEM of three technical replicates with the highest value on the y-axis reflecting the peptide’s maximum amount of possible deuterium uptake at 60 s. Black arrow points to H43, WT *Di*Cas7-11’s catalytic residue for pre-crRNA processing. Black circles represent apo Cas7-11 while orange circles represent *Di*Cas7-11 bound to crRNA.

### crRNA structurally rearranges regions near base-flipping centers

HDX-MS revealed that the domain-domain interfaces; Cas7.1/Cas7.2, Cas7.2/Cas7.3, and Cas7.3/Cas7.4 showed large FUDs upon crRNA binding (**Figure 4A**). The thumb-like β-hairpins of the Cas7.1-7.3 domains not only support the domain-domain interfaces through interaction with adjacent α1 helices, (e.g. Cas7.1 hairpin interacts with the Cas7.2 α1 helix) but also properly orient tgRNA for cleavage by interacting with crRNA and tgRNA nucleotides to induce base flips necessary for cleavage and sterically occlude these bases from interacting (3,15,18,21). Although the β-hairpin of Cas7.1 induces a base flip and interacts with the Cas7.2 α1 helix, it is not involved in cleavage because Cas7-11 has a shorter tgRNA than other type III effectors (3,18,33) and no conserved catalytic Asp in the Cas7.1 catalytic loop (16,18,34). The thumb-like β-hairpins of Cas7.1, covered by peptides 171-178 and 179-189 (**Figure 4D, shown in teal**) showed marked decreases in exchange upon crRNA binding (FUDs of -58.3% and -40.8% respectively). Thumb-like β-hairpins of Cas7.2 and Cas7.3, also showed decreases in deuterium uptake (FUD for residues 487-516 = -47.0%, residues 517-525 = -54.2%, residues 731-739 = -29.3%, residues 740-771 = -38.6%) (**Figure 4B and C**). The catalytic loops of Cas7.2 and Cas7.3, represented by peptides covering residues 411-423 (**Figure 4D**) and 634-657 (**Figure 4C**), also showed marked decreases in exchange (FUDs of -40.7% and -56.9%) upon crRNA binding. However, part of the Cas7.2 catalytic loop (residues 424-430), containing the active site residue, D429, showed no change (**Supplementary Figure S4B**), suggesting that structuring only occurs further down the catalytic loop (residues 411-423). Smaller peptides comprising the Cas7.3 catalytic loop residues 634-646 and 647-657 (which includes the Cas7.3 active site residue, D654) also showed strong protection (FUDs of -64.1% and -40.5%) showing that the entire Cas7.3 catalytic loop undergoes structuring (**Supplementary Figure S4A**). The Cas7.1-Cas7.4 α1 helices showed lower initial deuterium uptake in the apo state (**Figure 4A-D, Supplementary Figure S4A and S4C**) and lower FUDs compared to those seen in the β-hairpins and catalytic loops with the highest FUD, -20.4%, mapping to residues 431-457 of the Cas7.2 α1 helix. Some regions of the α1 helices showed almost no deuterium uptake before and after crRNA binding, suggesting that the α1 helices were well structured prior to crRNA binding (**Figures 4A-D, Supplementary Figure S4A and S4C**). Residues I504/V682 of Cas7.3 and M953/K1489, previously shown to “sandwich” the flipped 4^th^ and 10^th^ crRNA bases in the ternary complex structure (PDB:7WAH) (18) were contained in residues 487-516, 674-687, 1471-1497, and 937-965 that showed decreases in deuterium uptake upon crRNA binding (**Figure 4B and C and Supplementary Figure S4C**). Several other nearby regions covered by residues 680-687 and 1471-1477 with decreased deuterium uptake are also shown with their corresponding uptake plots in (**Supplementary Figure S4A and S4C**).

**Figure 4.**
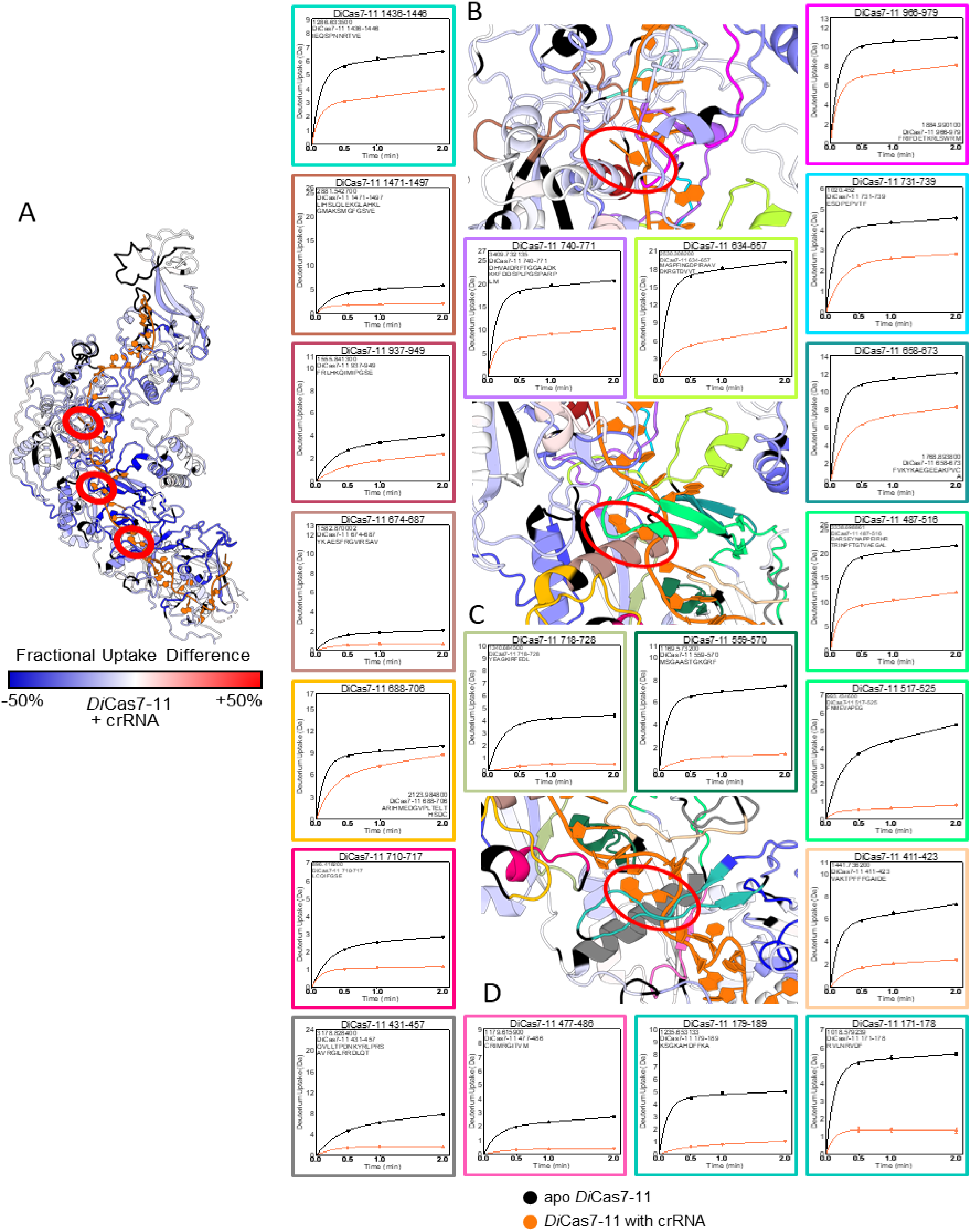
HDX-MS reveals that crRNA-binding induces protection of domain interfaces and catalytic sites. (**A**) FUD upon crRNA binding at 60 s deuterium incubation was mapped to *Di*Cas7-11-crRNA structure (PDB:7YN9). Flipped crRNA bases are circled in red. (**B**) Zoomed inset at the Cas7.3/Cas7.4 interface, colored by FUD and corresponding uptake plots, where Cas7.3 mediated target cleavage occurs. The flipped crRNA base is circled in red. (**C**) Zoomed inset at the Cas7.2/Cas7.3 interface where Cas7.2 mediated target cleavage occurs. (**D**) Zoomed inset at the Cas7.1/Cas7.2 interface.

### crRNA binding primes Cas11 domain for target cleavage

Biochemical evidence from Cas11 domain deletions as well as single point mutations indicate that Cas11 is necessary for tgRNA cleavage (18,20,35,36). It is thought that Cas11 properly orients the target RNA (18,21,35) while the β-hairpins flip the crRNA and tgRNA bases and keep them away from each other (3,18,21,37). Structural studies have shown that several Cas11 side chains interact with the tgRNA phosphate backbone and bases near cleavage sites, specifically the 3^rd^ and 9^th^ bases (18,20,35,36). Cas11 residues 274-284 and 285-298 showed -16.8% and -26.7% FUD respectively (**Figure 5A**). Residues 329-335 showed very low uptake values and slower deuterium uptake kinetics, and crRNA binding reduced the deuterium uptake to 0 (**Figure 5A**).

**Figure 5.**
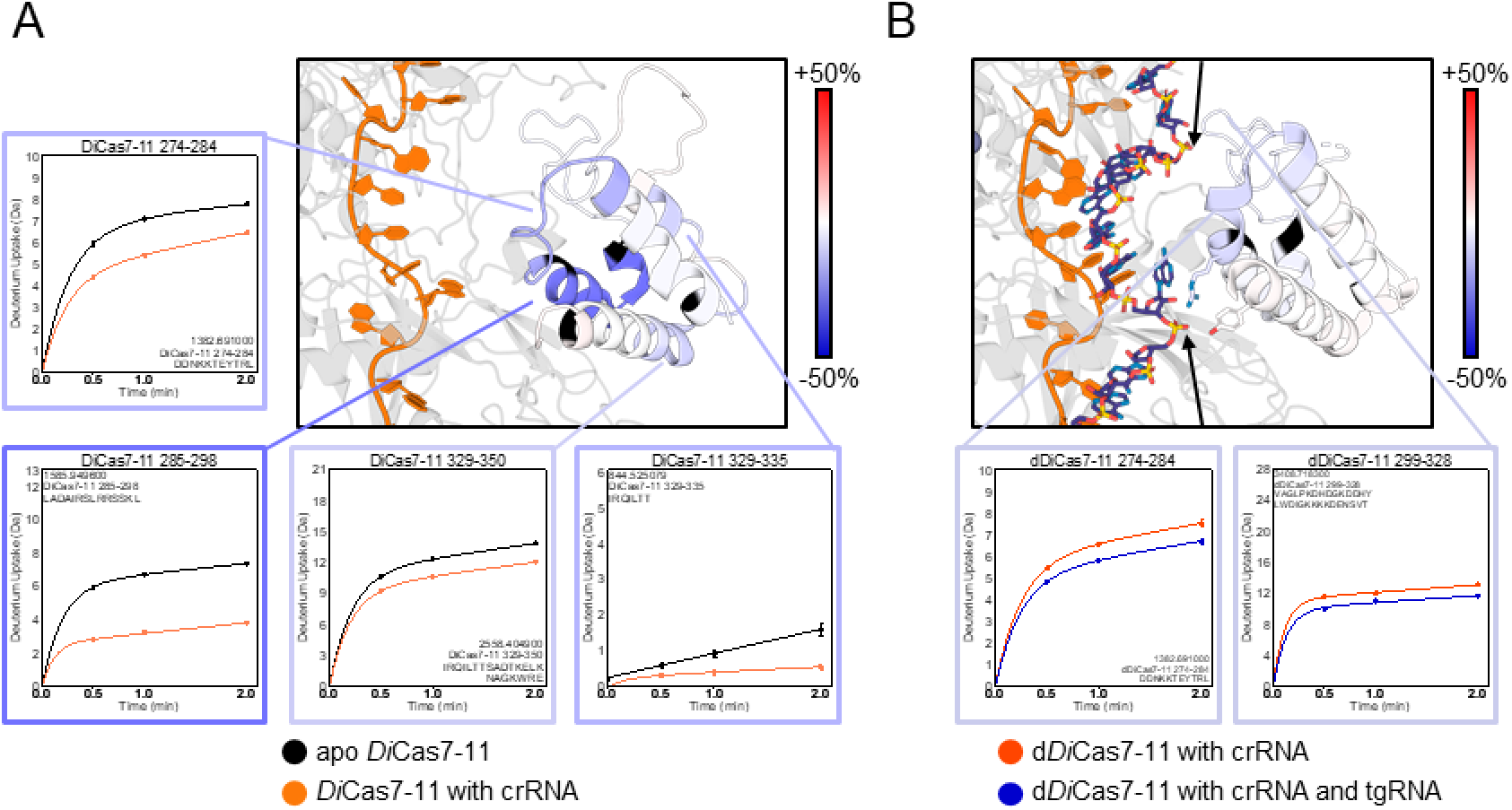
Structural changes in Cas11 domain from crRNA-binding in Cas11 domain but not tgRNA-binding. (**A**) FUD of crRNA-binding to WT *Di*Cas7-11 mapped to PDB:7YN9. Cas11 is colored by FUD with associated HDX uptake plots, while the rest of the protein is shown in grey. Lines from uptake plots point to their respective location on the structure. (**B**) FUD of tgRNA-binding to d*Di*Cas7-11-crRNA mapped to PDB:7WAH. Residues previously shown to be essential for cleavage activity in *Sb*Cas7-11 are shown as sticks. Scissile phosphates are marked by black arrows.

Target RNA binding marginally reduced exchange in residues of the Cas11 domain (**Figure 5B**). Whereas residues 274-284 showed a significant reduction in exchange upon crRNA binding (**Figure 5A**) only a slight additional reduction in exchange was observed upon tgRNA binding (FUD of -7.6%) (**Figure 5B**). Similarly, residues 285-298 and 299-328 showed little change upon tgRNA binding (**Figure 5B and Supplementary Figure S5**). Additional HDX uptake plots that show no changes upon crRNA/tgRNA interaction in other parts of Cas11 are shown in **Supplementary Figure S5**. HDX data suggests that the majority of structural ordering in Cas11 occurs prior to tgRNA binding with smaller, localized changes occurring upon tgRNA engagement.

### Target RNA binding induces localized changes near the active site residues responsible for target positioning for cleavage

Structural comparison of the binary (PDB:7YN9) and ternary complex (PDB:7YNA, 7WAH) suggest nearly identical structures except for a translation of Cas11 and the INS domain. Our HDX-MS results showed decreases in deuterium uptake localized to residues 424-430 and 634-657, which cover the inactivated catalytic residues D429A of Cas7.2 and D654A of Cas7.3, upon tgRNA binding (**Figure 6A and B**). Residues 634-646 showed no difference upon tgRNA binding, indicating that decrease in exchange upon tgRNA binding is localized to residues 647-657 (**Figure 6A**). Residues 411-423 representing the catalytic loop directly adjacent to the Cas7.2 active site, showed increased deuterium uptake upon tgRNA binding (**Figure 6B**). Adjacent increased and decreased HDX suggests a conformational rearrangement that may be involved in positioning of D429, although similar behavior was not seen near the Cas7.3 active site (**Figure 6A**). Another observation that suggests tgRNA induces conformational flexibility, was the bimodal deuterium envelope in the mass spectra for the peptide corresponding to residues 634-657 (**Figure 6C**). Bimodal deuterium exchange envelopes suggest the presence of two conformers, one of which is exchanging substantially more deuterium than the other (38–40). We interpret this to mean that tgRNA binding induces a slow opening of residues 634-657, relative to the deuterium incubation time scale. This behavior is also seen in residues 411-423, although to a lesser extent (**Figure 6D**).

**Figure 6.**
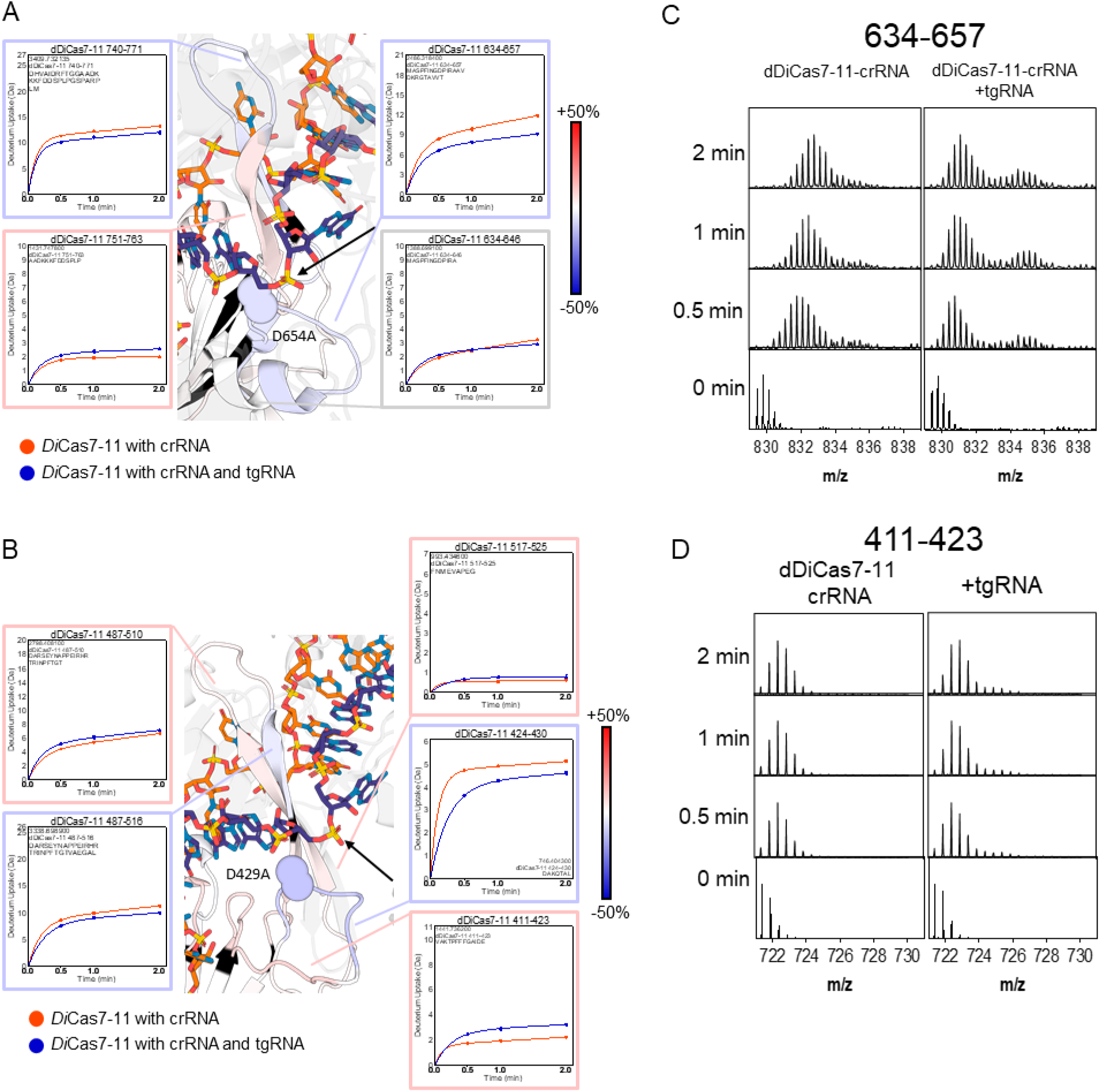
tgRNA-induced changes in d*Di*Cas7-11 active sites. (**A, B**) Zoomed-inset showing thumb-like β-hairpins and catalytic loops of Cas7.3 and Cas7.2 domains in a d*Di*Cas7-11-crRNA-tgRNA structure (PDB:7WAH). FUD of tgRNA binding was mapped to the structure with regions not showing significant FUD in grey. Orange-red circles represent d*Di*Cas7-11-crRNA while blue circles represent d*Di*Cas7-11-crRNA-tgRNA. Black arrows point to tgRNA scissile phosphates adjacent to flipped bases. (**C, D**) Raw mass spectra showing formation of a bimodal mass envelope distribution within catalytic loops of Cas7.3 and Cas7.2 domains upon tgRNA binding, suggesting two interconverting conformations with significantly different deuterium uptake.

Nearly a decade of type III effector characterization has implicated the role of thumb-like β-hairpins of Cas7 in flipping bases of the crRNA and tgRNA (15–18,21). Thus, it was surprising to see almost no difference in exchange within the β-hairpins of either Cas7.2 and Cas7.3 (**Figure 6A and B**) upon tgRNA binding. Comparison of residues 487-510 and 487-516 indicate a switch between a slight increase in FUD and a slight decrease in FUD upon tgRNA binding, respectively (**Figure 6B**). Thus, residues 511-516 in the Cas7.2 β-hairpin (**Figure 6B**, in light blue) may undergo some protection from interaction with the flipped 4^th^ base of tgRNA. Similarly, tgRNA binding leads to a FUD increase in residues 751-763 while a decrease is observed in residues 740-771 (**Figure 6A**). Since these two peptides partially overlap, these results imply a decrease in either 740-751, 764-771, or both.

### HDX-MS reveals structural flexibility of the INS domain in response to different RNAs

Independent studies have shown that deletion of the INS domain (*Di*Cas7-11Δ979-1293)(18,19) does not significantly affect RNA cleavage or knockdown, although one study showed an enhanced cleavage rate (21). Moreover, the structure of the INS domain varies significantly across orthologs (41) and is even required for target cleavage in *Sb*Cas7-11 (36). We also observed that deletion of the INS had no effect on catalytic activity (**Figure 7A**). EMSAs of WT *Di*Cas7-11-crRNA and *Di*Cas7-11(ΔINS)-crRNA showed that binding to tgRNA was also unaffected (**Figure 7B and C**). While an uptake plot of residues 1025-1037 (**Figure 7D**) indicated a decrease in exchange upon crRNA binding, further analysis of the mass spectra showed that this decrease resulted from the appearance of bimodal mass envelopes (**Figure 7E**), indicating two interconverting conformational states. HDX-MS analysis of the INS domain showed that bimodal mass spectral envelopes were observed across most of the INS domain (**Figure S6**) and mapped onto the *Di*Cas7-11-crRNA structure (PDB:7YN9) (**Figure 7F**). We also note that centroids of the higher m/z state in all analyzed peptides had the same m/z as the centroids observed for the apo state. These data can best be interpreted to indicate that crRNA binding induces an interconversion between a crRNA-interacting conformation (lower m/z state) and a second conformational state that is not interacting with the crRNA (higher m/z state).

**Figure 7.**
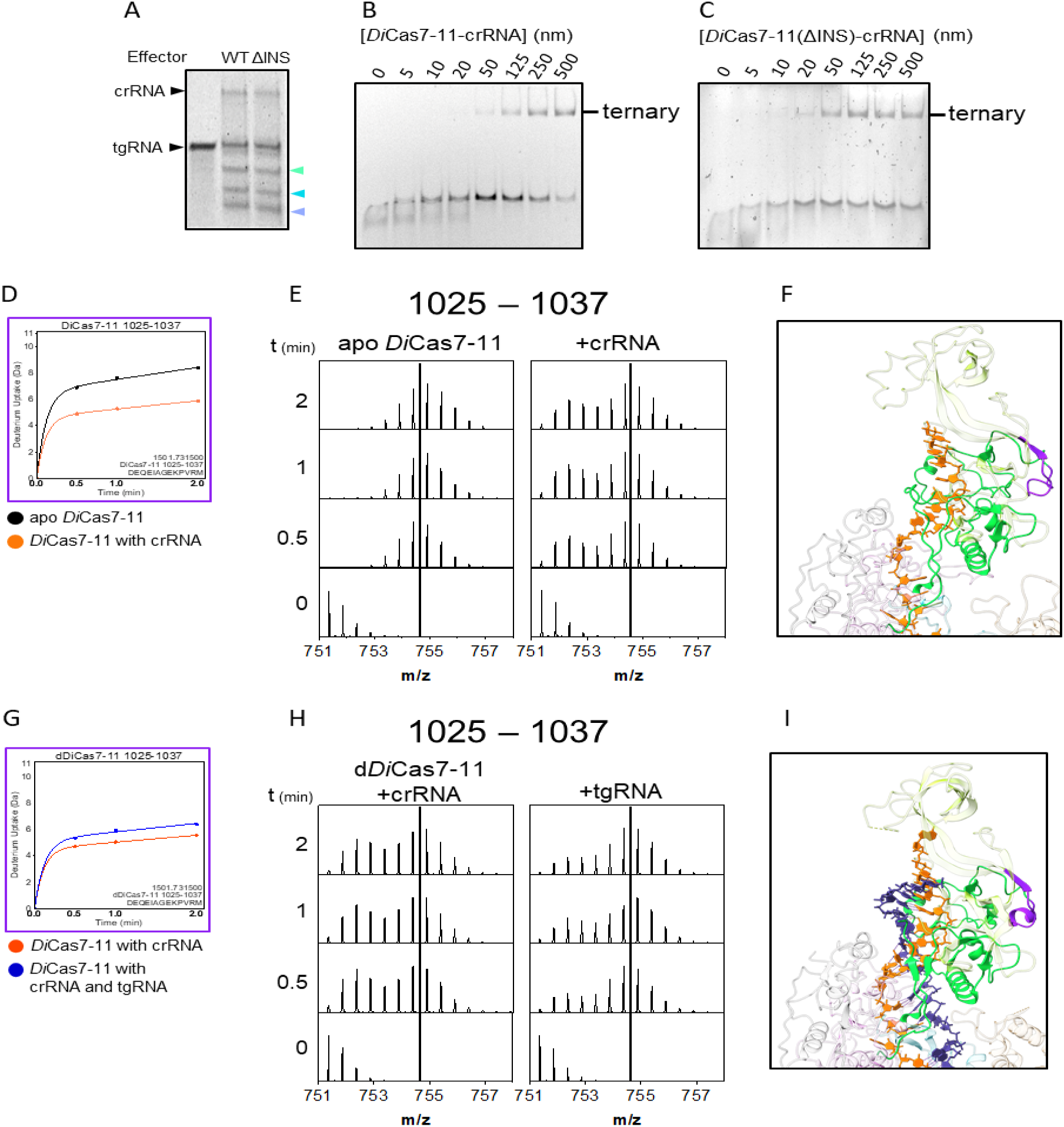
The INS domain is conformationally heterogeneous. (**A**) *in vitro* cleavage assay shows that INS deletion does not affect tgRNA cleavage. Target products were marked by their respective Cas7.2 and Cas7.3 domain colors. The lowest cleavage product was from Cas7.1 processing. (**B, C**) EMSAs of WT *Di*Cas7-11-crRNA and ΔINS *Di*Cas7-11-crRNA show similar tgRNA binding. (**D**) Uptake plot of residues 1025-1037 showing effects of crRNA binding. (**E**) Raw mass spectra of residues 1025-1037 showing formation of a bimodal mass envelope upon crRNA binding, indicating interconversion between two different conformations. Black lines through the y-axis show that the centroids have equal m/z values, suggesting that the associated conformations have a similar environment across states. (**F**) Structure of INS domain (PDB:7YN9) with residues 1025-1037 in purple and other residues exhibiting bimodal mass envelopes upon crRNA binding in green. The rest of the structure is transparent for ease of viewing. (**G**) Uptake plot of residues 1025-1037 showing effects of tgRNA binding. (**H**) Raw mass spectra of residues 1025-1037 showing an increase in the more exchanging conformation upon tgRNA binding, suggesting a more open INS domain. (**I**) Structure of INS domain (PDB:7WAH) with residues 1025-1037 in purple and other residues exhibiting bimodal mass envelopes upon crRNA binding in green.

Subsequent tgRNA binding to d*Di*Cas7-11-crRNA yielded a slight increase in deuterium uptake across the INS domain (**Figure 2C**) as represented by the peptide corresponding to residues 1025-1037 (**Figure 7G**). d*Di*Cas7-11-crRNA showed similar bimodal distributions for the INS domain as compared to the WT *Di*Cas7-11-crRNA. Subsequent tgRNA binding led to a decrease in the proportion of the less solvent exposed state (lower m/z) (**Figure 7H**). This effect was consistently observed across all mass spectra with bimodal distributions (**Supplementary Figure S7**) observed in the crRNA binding experiment and the regions of the INS which show this behavior are mapped onto the ternary complex structure PDB: 7WAH (**Figure 7I**). The more solvent exposed state had a centroid with similar m/z to the centroid observed in the apo WT state (**Figure 7E and Supplementary Figure S6 and S7**). Taken together, we interpret these results to mean that crRNA binding gives rise to an equilibrium between two states within the INS domain and that tgRNA binding shifts the equilibrium towards a conformation that is not interacting with the crRNA or the tgRNA.

### *Di*Cas7-11 does not process pre-crRNA on its own

It is not clear from previous work whether *Di*Cas7-11 can completely process a CRISPR array to a mature crRNA (11,18,25). To assess this, we compared the *Di*Cas7-11 processing activity of a single array-like transcript containing the direct repeat on both 5’ and 3’ ends of a 22-nt spacer versus the more commonly used 3’-less single array transcript with a direct repeat only on the 5’ end of a 22-nt spacer (11,18,19,21,42) (**Figure 8A-C**). While *Di*Cas7-11 processes a pre-crRNA with only a 5’ DR to the mature 37-nt crRNA, henceforth referred to as crRNA_37_, *Di*Cas7-11 processes the single array-like transcript into a partially mature crRNA, (referred to as crRNA_57_), that contains the mature 15-nt 5’-tag 5’ to the spacer and an additional 3’ 20-nt stem loop (**Figure 8C**). *In vitro* cleavage assays with *Di*Cas7-11-crRNA_57_ revealed reduced target cleavage (**Figure 8D**). Whereas tgRNA binding to *Di*Cas7-11-crRNA_37_ yielded only the highest band which we presume to be the active species (**Figure 8E**), *Di*Cas7-11-crRNA_57_ binding to tgRNA yielded multiple species with a lower population of the highest band. INS deletion mutants yielded a similar pattern (**Figure 8E**).

**Figure 8.**
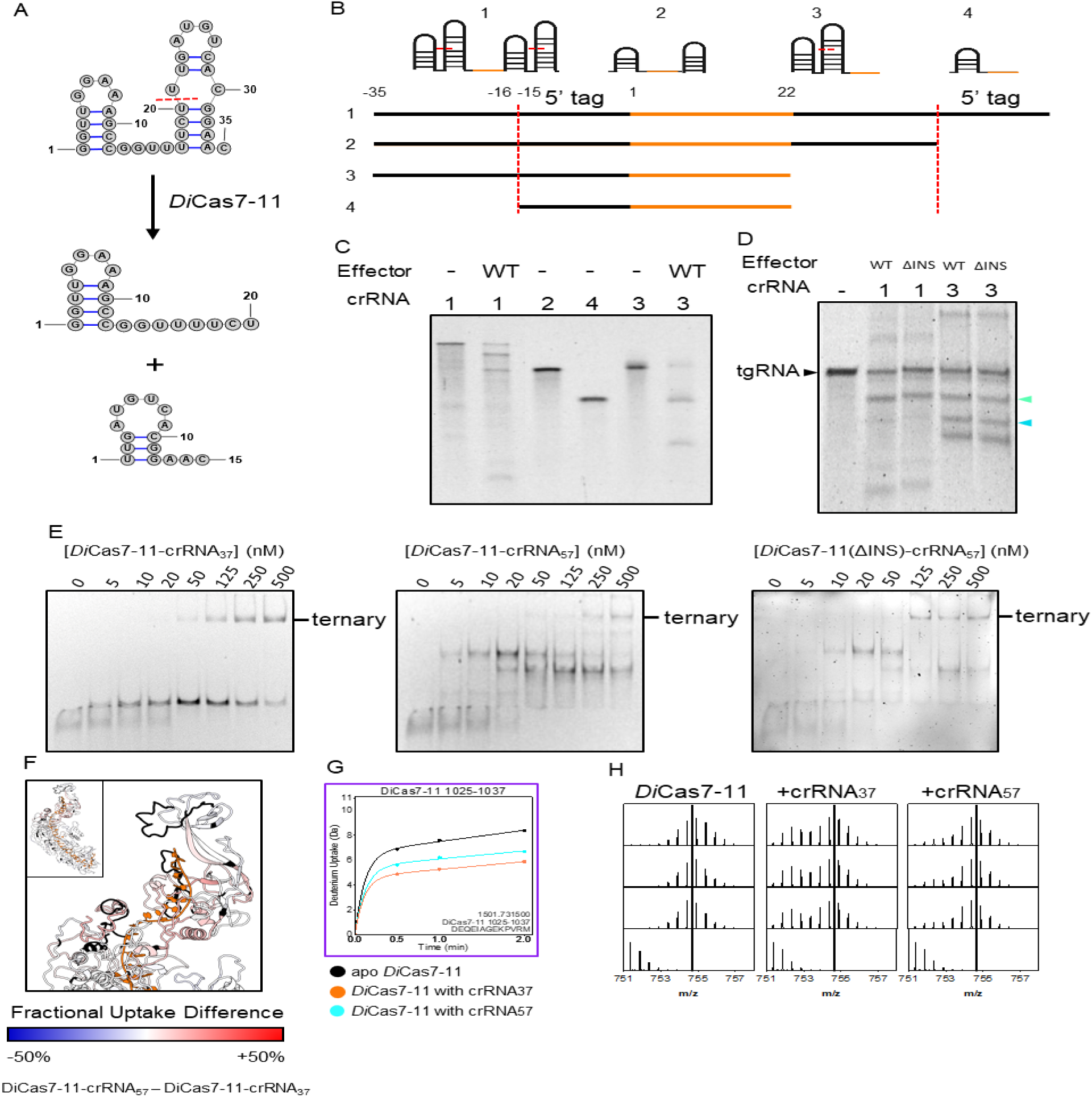
*in vitro Di*Cas7-11 pre-crRNA processing leads to a product that reduces tgRNA cleavage. (**A**) 35-nt *Di*Cas7-11 DR shown with secondary structure. Cas7.1 cleaves the DR at red dashed line, forming two products; a 15-nt 5’ tag recognized by the Cas7.1 domain and a 20-nt strand with stem-loop structure. (**B**) pre-crRNAs, 1 (single array) and 3 (5’DR-spacer), and crRNAs, 2 (crRNA_37_) and 4 (crRNA_57_), used for *in vitro* cleavage and binding assays are shown with Cas7.1 cleavage sites marked by red dashed lines. 3 and 4 are commonly used in processing assays. (**C**) *Di*Cas7-11 processing of various crRNAs. (**D**) Processing and target cleavage assays reveal the crRNA 2 (crRNA_57_) leads to lowered target cleavage. (**E**) tgRNA in the presence of increasing *Di*Cas7-11-crRNA_37_, *Di*Cas7-11-crRNA_57_, or *Di*Cas7-11(ΔINS)-crRNA_57_ reveal that crRNA_57_ leads to lowered amount of properly formed RNP. (**F**) Uptake plot of peptide 1025-1037. (**G**) FUD between *Di*Cas7-11-crRNA_57_ and *Di*Cas7-11-crRNA_37_ shows *Di*Cas7-11-crRNA_57_ has higher solvent accessibility near 5’ end of the tgRNA, where a stem-loop may be present. (**H**) Raw mass spectra of uptake plot showing varied distributions upon binding different crRNAs, suggesting a more open INS domain in *Di*Cas7-11-crRNA_57_ relative to *Di*Cas7-11-crRNA_37_.

Because our results indicated that *Di*Cas7-11 cannot fully process CRISPR RNA arrays on its own, and because RNase III was shown to process pre-crRNA of other Cas effectors (43), we compared the pre-crRNA processing ability of RNase III to that of *Di*Cas7-11. Our results show that RNase III is capable of processing the array, although the cleavage product had a slightly different size than crRNA_37_ (**Supplementary Figure S8**).

HDX-MS allowed for comparison of solvent accessibility between the WT *Di*Cas7-11 complex with the mature crRNA_37_ against the binary complex with the partially mature crRNA_57._ The additional 3’ 20-nt stem loop would be positioned somewhere after the last resolvable nucleotide at the 3’ region where the INS domain is located (PDB: 7YN9,7WAH, 8D1V (19)). Higher solvent accessibility throughout the INS domain was observed in the crRNA_57_ binary complex represented by residues 1025-1037 (**Figure 8F and 8G**). Mass spectra (**Figure 8H and Supplementary Figure S6**) revealed bimodal deuterium envelopes with less of the lower m/z envelope, reminiscent of those induced by tgRNA in the same peptides in the *Di*Cas7-11-crRNA_37_ sample. The *in vitro* processing and cleavage assays not only showed that *Di*Cas7-11 cannot process pre-crRNA itself but also that its processing product leads to reduced tgRNA cleavage. Taken together, binding assays and HDX-MS lead us to believe that the crRNA_57_ does not form an active ternary complex as effectively as crRNA_37_ which is reflected by a more open *Di*Cas7-11 INS domain.

### crRNA induces structural changes in TPR-CHAT interaction motifs

While our study focuses on the RNA-guided nuclease activity of *Di*Cas7-11, the data may shed light on how the RNA-guided protease Craspase complex forms. TPR-CHAT, or Csx29, as part of the Cas7-11 CRISPR loci, binds Cas7-11-crRNA to form the Craspase complex which is involved in the protease-mediated immune response (20,21,42,44,45). HDX-MS identified some changes in the known TPR-CHAT interaction motifs (20, 46). TPR-CHAT binds to Cas7-11-crRNA exclusively through a protein-protein interaction comprised of the TPR-CHAT N-terminal domain (NTD) interacting with residues 367-401 in *Di*Cas7-11 L2 and with residues 1313-1341 in the Cas7.4-2 domain (21,46). Both regions remain unresolved in the electron density of either Cas7-11-crRNA binary (or Cas7-11-crRNA-tgRNA ternary structures. HDX of the apo and binary WT *Di*Cas7-11 showed no change in the solvent exposed residues 367-401 (**Figure 9A**). Residues 1313-1341 in *Di*Cas7-11 were covered by two peptides; residues 1306-1322 which showed a decrease in deuterium uptake upon crRNA binding, and residues 1325-1346 which showed an increase (**Figure 9B**). The observed pattern of increase and decrease suggest that this region undergoes a conformational change upon crRNA binding. It is possible that the conformational change caused by crRNA binding induces the TPR-CHAT binding-competent conformation. tgRNA binding to d*Di*Cas7-11 did not appear to elicit significant changes in either interface (**Supplementary Figure S9**).

**Figure 9.**
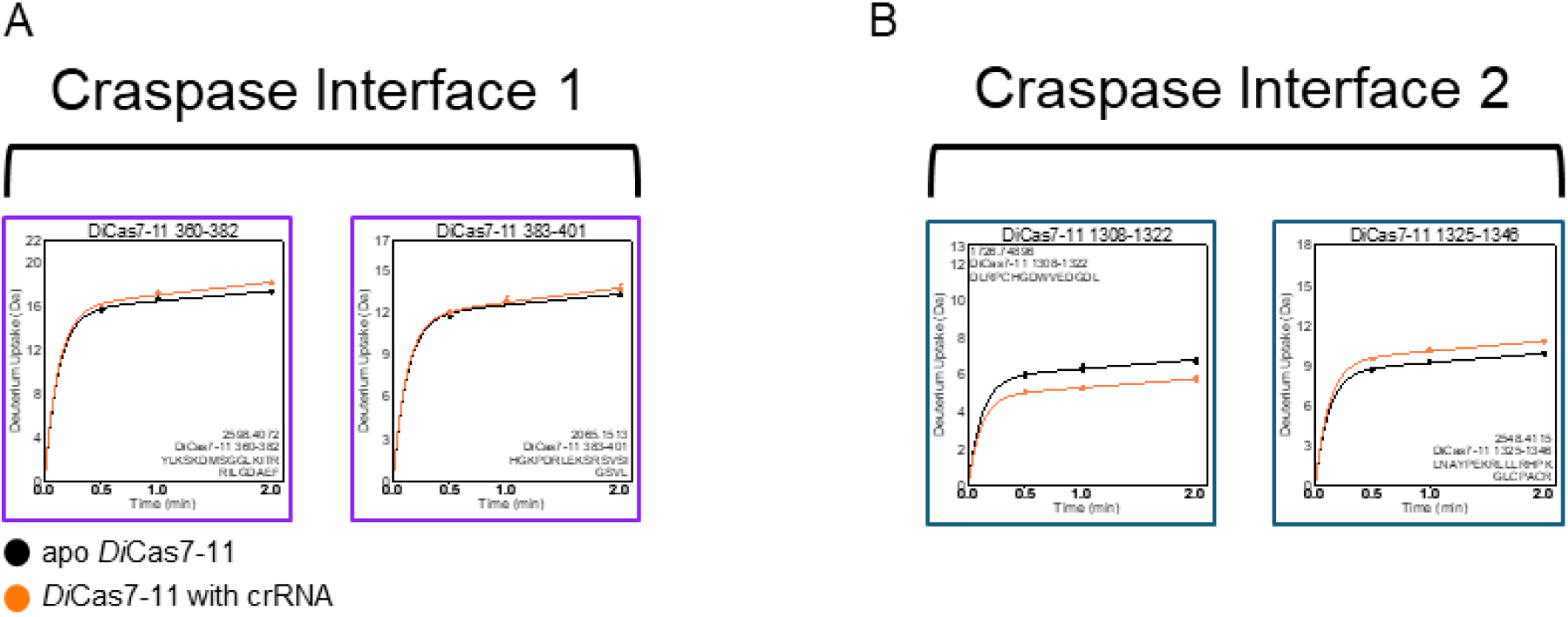
HDX-MS analysis of the regions of *Di*Cas7-11-crRNA responsible for TPR-CHAT binding (**A**) Craspase interface 1 remains solvent exposed upon crRNA binding. (**B**) A decrease in exchange in residues 1308-1322 and an increase in exchange in residues 1325-1346 was observed, suggests a conformational change within Craspase interface 2.

## DISCUSSION

HDX-MS analysis of apo Cas7-11 provided critical information regarding what “holds” the four Cas7 domains together. The Cas7 domains are RRM folds that bind zinc, but also have sequence inserts that bind the crRNA, induce base flipping, and catalyze target cleavage. The four Cas7 domains form an overall crescent shape with the crRNA bound in the inner concave surface (18,19,21,47). HDX-MS revealed that the convex side of the crescent shaped Cas7-11, formed by the RRM folds consisting of the β1-α1-β2-β3-α2-β4 structure in each Cas7 domain had low deuterium exchange. This is likely to be at least partly due to the zinc-binding which stabilizes the fold and does not occur in the Csm/Cmr complexes (3,18). In contrast, the concave surface corresponding to the additional motifs inserted into the RRM folds were highly exchanging. Thus, the crRNA recognition motifs appear to fold upon binding to the crRNA but are held in position by the more rigidly folded sequential RRM domains.

Cas7-11 appears to have evolved from the linking of four separate Cas7 protein domains (11,31), so we wondered whether there were interactions between the Cas7 domains or whether they were more like beads-on-a-string in the apo state. The cryoEM structures of the binary and ternary RNA-bound Cas7-11 (PDB: 7YN9 and 7WAH) forms highlighted two interfaces; a primary domain composed of the β1-α1 and α2-β4 regions of a Cas7 domain (e.g. Cas7.2) interacting with a β3 strand of an adjacent Cas7 domain (Cas7.3) and a second interface in which each thumb-like β-hairpin extends from the previous Cas7 domain to the α1-helix of the next. HDX analysis of the apo state showed that some of these motifs were solvent exposed, specifically the β1-α1 of Cas7.1-Cas7.3 and α2-β4 of Cas7.2, indicating they are unlikely to form stable interfaces prior to crRNA binding. Instead, we find that the L3 linker (*cf*. **Figure 2A**) is surprisingly protected from exchange and appears to be what connects Cas7.2 to Cas7.3 through stabilizing hydrophobic interactions between α2 and L3. The Cas7.2-Cas7.1 interface may also be stabilized by an L3 interaction with the Cas7.1 α2 helix. A long loop in the CTE that extends from the Cas7.4 β4 strand and interacts with α2 from Cas7.3 is also protected from exchange and likely stabilizes the Cas7.3-Cas7.4 interface. The previously described α2-β4 of Cas7.3 to the β3 of Cas7.4 interface was also solvent protected, suggesting that this interaction may be present prior to crRNA binding (18). HDX-MS demonstrates that the RRM folds of *Di*Cas7-11 are well-structured and, along with linkers and the CTE, form domain-domain interfaces rather than a beads-on-a-string architecture in the apo state. In contrast, the sequences inserted into the RRM fold core sequences that are responsible for crRNA binding and cleavage are highly dynamic.

Large decreases in solvent exchange indicative of coupled folding and binding were observed along the crRNA-binding surface. The thumb-like β-hairpins that extend from Cas7.1 to Cas7.2, Cas7.2 to Cas7.3, and Cas7.3 to Cas7.4 all show dramatic decreases in exchange upon crRNA binding. These appear to solidify the domain-domain interactions and sandwich the flipped base in the crRNA against an adjacent domain’s α1-helix (e.g. Cas7.2 β-hairpin to Cas7.3 α1-helix) (18,19,21). Cas7.2 and Cas7.3 also have catalytic loops that become dramatically more ordered. The magnitude of deuterium uptake changes in the catalytic loops and β-hairpins (**Figure 4B-D**) suggest crRNA mediated structural ordering. Similarly, electron density for the thumb-like β-hairpin and catalytic loop in the x-ray structure of the apo state (PDB: 4QTS) of the Cas7-like Csm3 protein of the ancestral type III-A effectors became visible loops in the cryoEM structure of the crRNA-bound state (PDB: 6MUU) (15,48). Finally, regions of the processing site of Cas7.1 showed marked decreases in deuterium uptake upon crRNA binding. The β1 strand containing pre-crRNA processing catalytic residue H43 became ordered upon binding as did other important binding residues (18,21).

Target RNA binding to the Cas7 backbone showed small decreases in exchange localized to the catalytic loops. These decreases were much less pronounced than those seen upon crRNA binding, implying that crRNA-binding is primarily responsible for folding the disordered regions of Cas7-11. A concomitant increase in solvent accessibility in residues 411-423 and decrease in solvent accessibility in residues 424-430 of Cas7.2 (**Figure 4B**) suggests a rearrangement of the flexible catalytic loop, which may facilitate proper positioning of the catalytic residue. Flexibility of a different kind is seen in the catalytic loop of Cas7.3, residues 634-657, which has a bimodal deuterium uptake distribution suggesting two slowly interconverting states.

The marked decrease in exchange in both Cas11 α2 helix (residues 283-295) and the helical turn of the Cas7.3 catalytic loop upon crRNA-binding suggests formation of a new interface between Cas11 and the Cas7.3 domain (19,21). This new interaction may be facilitated by a nascent hydrophobic interaction between P643 and a crRNA base at position 5. The Cas11 α2 helix residue R283 which appears to be ordered upon crRNA binding has been implicated in positioning the scissile phosphate in structural (18,19,21,46) and mutagenesis studies (20,35). The lack of changes in deuterium uptake upon tgRNA binding throughout the Cas11 domain (**Figure 5B and Supplementary Figure S5**) is consistent with previous structural comparisons between the binary and ternary complexes that show only slight translations and rotations (21,35) in Cas11 upon tgRNA binding. Thus, coordination of scissile phosphates appears to be mediated by crRNA binding (**Figure 5B, black arrows**).

The literature is unclear as to whether *Di*Cas7-11 can process a mature crRNA from the CRISPR array transcript *in vitro* (10). Studies co-expressing *Di*Cas7-11 or *Sb*Cas7-11 with their respective CRISPR arrays in *E. coli* have shown that crRNAs are processed to their mature sizes, with 15-nt and 18-nt 5’ tags respectively followed by ∼20-nt spacers (11,12). Using a single array transcript, we showed that *Di*Cas7-11 leaves an additional 20-nt 3’ hairpin (as seen in crRNA_57_). This lack of complete processing was not known because most studies in the literature use a pre-crRNA with no 3’ DR and consequently no 3’ hairpin upon processing (crRNA_37_). The observation that *Di*Cas7-11 is unable to process pre-crRNA makes sense because its ortholog *Sb*Cas7-11 is also unable to process pre-crRNA (12,35,36,47,49) and type III-A/B complexes require other nucleases for crRNA processing (3). These results strongly suggest that for all Cas7-11 proteins an ancillary nuclease such as RNase III, from the host organism will be needed to cleave the DR 3’ to the spacer.

The INS domains of type III-E Cas effectors are highly variable and their function is unknown. The INS domains in *Di*Cas7-11 are only partly resolvable (PDB: 7Y9X, 8GS2) (21,32) or not at all visible (PDB: 8D1V, 8WMC, 8WMI, 8WM4, 8EEX) (19,42,46) in the available structures. No density for the INS domain of *Sb*Cas7-11 (*Sb*gRAMP) has been observed in any structural studies (20,35,36,47,49) suggesting that *Sb*Cas7-11 has a more flexible INS than *Di*Cas7-11. Our HDX data is consistent with the INS domain being highly flexible. Functionally, the INS domain of *Di*Cas7-11 can be deleted and the catalytic activity for tgRNA doesn’t change (18,19). On the other hand, charge reversal mutations in the INS domain (D982K/D988K/D1581R) appeared to increase RNA knockdown in human cells (13). It remains a complete mystery as to what the INS might be doing, if anything, in *Di*Cas7-11.

The 3’ 20-nt hairpin in crRNA_57_ would be positioned near the INS, so we probed whether the INS could sense the difference between this partially processed crRNA and the fully-processed crRNA_37._ *Di*Cas7-11-crRNA_37_ formed a single complex with tgRNA according to EMSA assays, whereas *Di*Cas7-11-crRNA_57_ formed several differently eluting species. HDX-MS revealed that the entire INS domain was interconverting between two different conformations upon crRNA binding, one that was more protected and one that was significantly less protected from exchange. More of the protected conformation was observed in *Di*Cas7-11-crRNA_37_ when compared to *Di*Cas7-11-crRNA_57_. The predominance of the less protected conformation suggests that the INS domain does not interact with the crRNA_57_ as well as with the crRNA_37_. The fact that *Di*Cas7-11-crRNA_57_ shows lower cleavage activity than *Di*Cas7-11-crRNA_37_ is consistent with the idea that only the fully-formed complex is catalytically active. Upon tgRNA binding to *Di*Cas7-11-crRNA_37,_ the INS conformational equilibrium shifts to the less protected state. In other words, the INS domain appears to be in a mostly open state in the ternary complex which could explain why INS domain deletion does not appear to impair tgRNA cleavage activity. The increased exposure of the INS upon tgRNA binding suggests that the INS is disengaged during target cleavage consistent with the observation of equal cleavage activity for *Di*Cas7-11(ΔINS)-crRNA vs. *Di*Cas7-11-crRNA. While the INS domain from the type III-Dv effector has been shown to “seed” tgRNA binding (24), *Di*Cas7-11 does not have the residues implicated in this function. Taken together with the HDX data, our results suggest that the *Di*Cas7-11 INS domain is vestigial.

Finally, it is thought that crRNA binding promotes TPR-CHAT binding. The HDX-MS results reveal a simultaneous decrease and increase in solvent accessibility within one of the disordered TPR-CHAT binding motifs of *Di*Cas7-11 upon crRNA binding. Such yin-yang HDX behavior is evidence of a conformational change that may properly orient the *Di*Cas7-11 binding region for TPR-CHAT binding.

Our results provide a first glimpse of an apo type III-E effector, revealing its conformationally flexible nature. Several key events arising from crRNA binding were pivotal in forming a functional protein; namely the folding of initially disordered catalytic loops and disordered β-hairpins, stronger protein interactions at domain-domain interfaces, and the folding of various regions surrounding the Cas7.1 processing site. This work expands on the growing body of literature on the protein dynamics of CRISPR Cas effectors and further supports a model in which the apo protein is disordered and forms a catalytically active structure upon crRNA binding. A lack of structural information on the apo state underscores the importance of using biophysical methods along with structural biology (50). As the first RNA-targeting single-protein effector without collateral cleavage (11), biophysical characterization Cas7-11 may prove informative for developing RNA editing tools and therapies (51,52).

## DATA AVAILABILITY

The raw HDXMS data files and analyzed data are available at massive.ucsd.edu dataset MSV000095504. Uptake plots may be generated from the state data excel file using the DECA program available at https://github.com/komiveslab/DECA

## SUPPLEMENTARY DATA

Supplementary Data are available online.

## AUTHOR CONTRIBUTIONS

C.P.L.: investigation, methodology, formal analysis, conceptualization, writing – original draft. H.L.: investigation. TW: investigation. D.J.B investigation, methodology, writing – review & editing. O.A.S.: supervision, resources. E.A.K. – supervision, resources, methodology, writing – original draft.

## ACKNOWLEDGEMENTS

We thank Steve Silletti for assistance and mentorship in operating the mass spectrometer.

## REFERENCES

1. Barrangou, R., Fremaux, C., Deveau, H., Richards, M., Boyaval, P., Moineau, S., Romero, D.A. and Horvath, P. (2007) CRISPR provides acquired resistance against viruses in prokaryotes. Science, 315, 1709–1712.

2. Hille, F., Richter, H., Wong, S.P., Bratovič, M., Ressel, S. and Charpentier, E. (2018) The Biology of CRISPR-Cas: Backward and Forward. Cell, 172, 1239–1259.

3. Liu, T.Y. and Doudna, J.A. (2020) Chemistry of Class 1 CRISPR-Cas effectors: Binding, editing, and regulation. J Biol Chem, 295, 14473–14487.

4. Liu, L., Li, X., Wang, J., Wang, M., Chen, P., Yin, M., Li, J., Sheng, G. and Wang, Y. (2017) Two Distant Catalytic Sites Are Responsible for C2c2 RNase Activities. Cell, 168, 121–134.e112.

5. Yan, W.X., Chong, S., Zhang, H., Makarova, K.S., Koonin, E.V., Cheng, D.R. and Scott, D.A. (2018) Cas13d Is a Compact RNA-Targeting Type VI CRISPR Effector Positively Modulated by a WYL-Domain-Containing Accessory Protein. . Mol. Cell, 70, 327–339.e325.

6. Yang, H. and Patel, D.J. (2024) Structures, mechanisms and applications of RNA-centric CRISPR-Cas13. Nat Chem Biol, 20, 673–688.

7. Kellner, M.J., Koob, J.G., Gootenberg, J.S., Abudayyeh, O.O. and Zhang, F. (2019) SHERLOCK: nucleic acid detection with CRISPR nucleases. Nat. Protoc., 14, 2986–3012.

8. Brogan, D.J., Chaverra-Rodriguez, D., Lin, C.P., Smidler, A.L., Yang, T., Alcantara, L.M., Antoshechkin, I., Liu, J., Raban, R.R., Belda-Ferre, P. et al. (2021) Development of a Rapid and Sensitive CasRx-Based Diagnostic Assay for SARS-CoV-2. ACS Sens, 6, 3957–3966.

9. Buchman, A.B., Brogan, D.J., Sun, R., Yang, T., Hsu, P.D. and Akbari, O.S. (2020) Programmable RNA Targeting Using CasRx in Flies. CRISPR J, 3, 164–176.

10. Wang, Q., Liu, X., Zhou, J., Yang, C., Wang, G., Tan, Y., Wu, Y., Zhang, S., Yi, K. and Kang, C. (2019) The CRISPR-Cas13a Gene-Editing System Induces Collateral Cleavage of RNA in Glioma Cells. Adv. Sci., 6, 1901299.

11. Özcan, A., Krajeski, R., Ioannidi, E., Lee, B., Gardner, A., Makarova, K.S., Koonin, E.V., Abudayyeh, O.O. and Gootenberg, J.S. (2021) Programmable RNA targeting with the single-protein CRISPR effector Cas7-11. Nature, 597, 720–725.

12. van Beljouw, S.P.B., Haagsma, A.C., Rodríguez-Molina, A., van den Berg, D.F., Vink, J.N.A. and Brouns, S.J.J. (2021) The gRAMP CRISPR-Cas effector is an RNA endonuclease complexed with a caspase-like peptidase. Science, 373, 1349–1353.

13. Schmitt-Ulms, C., Kayabolen, A., Manero-Carranza, M., Zhou, N., Donnelly, K., Nuccio, S.P., Kato, K., Nishimasu, H., Gootenberg, J.S. and Abudayyeh, O.O. (2024) Programmable RNA writing with trans-splicing. bioRxiv, 10.1101/2024.1101.1131.578223v578221.

14. Nemudraia, A., Nemudryi, A. and Wiedenheft, B. (2024) Repair of CRISPR-guided RNA breaks enables site-specific RNA excision in human cells. Science, 384, 808–814.

15. Jia, N., Mo, C.Y., Wang, C., Eng, E.T., Marraffini, L.A. and Patel, D.J. (2019) Type III-A CRISPR-Cas Csm Complexes: Assembly, Periodic RNA Cleavage, DNase Activity Regulation, and Autoimmunity. . Mol. Cell, 73, 264–277.e265.

16. Taylor, D.W., Zhu, Y., Staals, R.H.J., Kornfeld, J.E., Shinkai, A., van der Oost, J., Nogales, E. and Doudna, J.A. (2015) Structural biology. Structures of the CRISPR-Cmr complex reveal mode of RNA target positioning. Science, 348, 581–585.

17. Osawa, T., Inanaga, H., Sato, C. and Numata, T. (2015) Crystal structure of the CRISPR-Cas RNA silencing Cmr complex bound to a target analog. . Mol. Cell, 58, 418– 430.

18. Kato, K., Zhou, W., Okazaki, S., Isayama, Y., Nishizawa, T., Gootenberg, J.S., Abudayyeh, O.O. and Nishimasu, H. (2022) Structure and engineering of the type III-E CRISPR-Cas7-11 effector complex. Cell, 185, 2324–2337.e2316.

19. Goswami, H.N., Rai, J., Das, A. and Li, H. (2022) Molecular mechanism of active Cas7-11 in processing CRISPR RNA and interfering target RNA. Elife, 11, e81678.

20. Hu, C., van Beljouw, S.P.B., Nam, K.H., Schuler, G., Ding, F., Cui, Y., Rodríguez-Molina, A., Haagsma, A.C., Valk, M., Pabst, M., et al. (2022) Craspase is a CRISPR RNA-guided, RNA-activated protease. Science, 377, 1278–1285.

21. Huo, Y., Zhao, H., Dong, Q. and Jiang, T. (2023) Cryo-EM structure and protease activity of the type III-E CRISPR-Cas effector. Nat Microbiol, 8, 522–532.

22. Behler, J., Sharma, K., Reimann, V., Wilde, A., Urlaub, H. and Hess, W.R. (2018) The host-encoded RNase E endonuclease as the crRNA maturation enzyme in a CRISPR-Cas subtype III-Bv system. Nat Microbiol, 3, 367–377.

23. Hatoum-Aslan, A., Samai, P., Maniv, I., Jiang, W. and Marraffini, L.A. (2013) A ruler protein in a complex for antiviral defense determines the length of small interfering CRISPR RNAs. J Biol Chem, 288, 27888–27897.

24. Schwartz, E.A., Bravo, J.P.K., Ahsan, M., Macias, L.A., McCafferty, C.L., Dangerfield, T.L., Walker, J.N., Brodbelt, J.S., Palermo, G., Fineran, P.C. et al. (2024) RNA targeting and cleavage by the type III-Dv CRISPR effector complex. Nat. Commun, 15, 3324.

25. Brogan, D.J., Benetta, E.D., Wang, T., Lin, C.P., Chen, F., Li, H., Lin, C., Komives, E.A. and Akbari, O.S. (2024) Synthetic type III-E CRISPR-Cas effectors for programmable RNA-targeting. bioRxiv, 10.1101/2024.1102.1123.581838.

26. Malakhov, M.P., Mattern, M.R., Malakhova, O.A., Drinker, M., Weeks, S.D. and Butt, T.R. (2004) SUMO fusions and SUMO-specific protease for efficient expression and purification of proteins. J Struct Funct Genomics, 5, 75–86.

27. Lumpkin, R.J., Ahmad, A.S., Blake, R., Condon, C.J. and Komives, E.A. (2021) The Mechanism of NEDD8 Activation of CUL5 Ubiquitin E3 Ligases. . Mol Cell Proteomics, 20, 100019.

28. Lumpkin, R.J. and Komives, E.A. (2019) DECA, A Comprehensive, Automatic Post-processing Program for HDX-MS Data. Mol Cell Proteomics, 18, 2516–2523.

29. Masson, G.R., Burke, J.E., Ahn, N.G., Anand, G.S., Borchers, C., Brier, S., Bou-Assaf, G.M., Engen, J.R., Englander, S.W., Faber, J. et al. (2019) Recommendations for performing, interpreting and reporting hydrogen deuterium exchange mass spectrometry (HDX-MS) experiments. Nat Methods, 16, 595–602.

30. Komives, E.A. (2023) Dynamic allostery in thrombin-a review. Front Mol Biosci, 10, 1200465.

31. Makarova, K.S., Wolf, Y.I., Iranzo, J., Shmakov, S.A., Alkhnbashi, O.S., Brouns, S.J.J., Charpentier, E., Cheng, D., Haft, D.H., Horvath, P. et al. (2020) Evolutionary classification of CRISPR-Cas systems: a burst of class 2 and derived variants. Nat Rev Microbiol, 18, 67–83.

32. Kato, K., Okazaki, S., Schmitt-Ulms, C., Jiang, K., Zhou, W., Ishikawa, J., Isayama, Y., Adachi, S., Nishizawa, T., Makarova, K.S. et al. (2022) RNA-triggered protein cleavage and cell growth arrest by the type III-E CRISPR nuclease-protease. Science, 378, 882– 889.

33. Kolesnik, M.V., Fedorova, I., Karneyeva, K.A., Artamonova, D.N. and Severinov, K.V. (2021) Type III CRISPR-Cas Systems: Deciphering the Most Complex Prokaryotic Immune System. Biochemistry (Mosc), 86, 1301–1314.

34. Benda, C., Ebert, J., Scheltema, R.A., Schiller, H.B., Baumgärtner, M., Bonneau, F., Mann, M. and Conti, E. (2014) Structural model of a CRISPR RNA-silencing complex reveals the RNA-target cleavage activity in Cmr4. Mol Cell, 56, 43–54.

35. Yu, G., Wang, X., Zhang, Y., An, Q., Wen, Y., Li, X., Yin, H., Deng, Z. and Zhang, H. (2022) Structure and function of a bacterial type III-E CRISPR-Cas7-11 complex. Nat Microbiol, 7, 2078–2088.

36. Liu, X., Zhang, L., Wang, H., Xiu, Y., Huang, L., Gao, Z., Li, N., Li, F., Xiong, W., Gao, T. et al. (2022) Target RNA activates the protease activity of Craspase to confer antiviral defense. Mol Cell, 82, 4503–4518.

37. Nishimasu, H. and Nureki, O. (2017) Structures and mechanisms of CRISPR RNA-guided effector nucleases. Curr Opin Struct Biol, 43, 68–78.

38. Konermann, L., Tong, X. and Pan, Y. (2008) Protein structure and dynamics studied by mass spectrometry: H/D exchange, hydroxyl radical labeling, and related approaches. . J Mass Spectrom, 43, 1021–1036.

39. James, E.I., Murphree, T.A., Vorauer, C., Engen, J.R. and Guttman, M. (2022) Advances in Hydrogen/Deuterium Exchange Mass Spectrometry and the Pursuit of Challenging Biological Systems. Chem Rev, 122, 7562–7623.

40. Torres-Paris, C., Song, H.J., Engelberger, F., Ramírez-Sarmiento, C.A. and Komives, E.A. (2024) The Light Chain Allosterically Enhances the Protease Activity of Murine Urokinase-Type Plasminogen Activator. Biochemistry, 63, 1434–1444.

41. Ekundayo, B., Torre, D., Beckert, B., Nazarov, S., Myasnikov, A., Stahlberg, H. and Ni, D. (2023) Structural insights into the regulation of Cas7-11 by TPR-CHAT. Nat Struct Mol Biol, 30, 135–139.

42. Strecker, J., Demircioglu, F.E., Li, D., Faure, G., Wilkinson, M.E., Gootenberg, J.S., Abudayyeh, O.O., Nishimasu, H., Macrae, R.K. and Zhang, F. (2022) RNA-activated protein cleavage with a CRISPR-associated endopeptidase. Science, 378, 874–881.

43. Deltcheva, E., Chylinski, K., Sharma, C.M., Gonzales, K., Chao, Y., Pirzada, Z.A., Eckert, M.R., Vogel, J. and Charpentier, E. (2011) CRISPR RNA maturation by trans-encoded small RNA and host factor RNase III. Nature, 471, 602–607.

44. Stella, G. and Marraffini, L. (2024) Type III CRISPR-Cas: beyond the Cas10 effector complex. Trends Biochem Sci, 49, 28–37.

45. Burgess, D.J. (2023) New cuts for CRISPR effectors. Nat Rev Genet, 24, 71.

46. Hong, T., Luo, Q., Ma, H., Wang, X., Li, X., Shen, C., Pang, J., Wang, Y., Chen, Y., Zhang, C. et al. (2024) Structural basis of negative regulation of CRISPR-Cas7-11 by TPR-CHAT. Signal Transduct Target Ther, 9, 111.

47. Cui, N., Zhang, J.-T., Li, Z., Liu, X.-Y., Wang, C., Huang, H. and Jia, N. (2022) Structural basis for the non-self RNA-activated protease activity of the type III-E CRISPR nuclease-protease Craspase. . Nat Commun, 13, 7549.

48. Numata, T., Inanaga, H., Sato, C. and Osawa, T. (2015) Crystal structure of the Csm3-Csm4 subcomplex in the type III-A CRISPR-Cas interference complex. J Mol Biol, 427, 259–273.

49. Wang, S., Guo, M., Zhu, Y., Lin, Z. and Huang, Z. (2022) Cryo-EM structure of the type III-E CRISPR-Cas effector gRAMP in complex with TPR-CHAT. Cell Res, 32, 1128– 1131.

50. Engen, J.R. and Komives, E.A. (2020) Complementarity of Hydrogen/Deuterium Exchange Mass Spectrometry and Cryo-Electron Microscopy. Trends Biochem. Sci., 45, 906–918.

51. Koonin, E.V., Gootenberg, J.S. and Abudayyeh, O.O. (2023) Discovery of Diverse CRISPR-Cas Systems and Expansion of the Genome Engineering Toolbox. Biochemistry, 62, 3465–3487.

52. Catchpole, R.J. and Terns, M.P. (2021) New Type III CRISPR variant and programmable RNA targeting tool: Oh, thank heaven for Cas7-11. Mol Cell, 81, 4354– 4356.

