## supplemental figures for "CRISPR-RNA binding drives structural ordering that primes Cas7-11 for target cleavage"

Table S1. RNA oligonucleotides used for this study

| | RNA | RNA sequence (5' --> 3') Direct repeat in red | Production | Expt | Figure(s) |
|---|---|---|---|---|---|
| 1 | Di-crRNA-array | gguuggaaagccgguuuucuuugaugucacggaaccugguagaauuucugugguaacgguuggaaagccgguuuucuuugaugucacggaac | Ordered from IDT | cleavage | Figure 8 |
| 2 | Di-crRNA-22nt_spacer | GGGgguuggaaagccgguuuucuuugaugucacggaaccugguagaauuucugugguaac | *in vitro* transcription | cleavage | Figure 7,8 |
| 3 | Di-crRNA-22nt_spacer | gguuggaaagccgguuuucuuugaugucacggaaccugguagaauuucugugguaac | Ordered from IDT | HDX, EMSA | Figure 2-9 |
| 4 | Di-crRNA-22nt_spacer-20DR | gguuggaaagccgguuuucuuugaugucacggaaccugguagaauuucugugguaacgguuggaaagccgguuuucu | Ordered from IDT | HDX, EMSA | Figure 8 |
| 5 | FAM-labeled target RNA | 6-FAM-AAUUUUACUAUUAGUGUUACCACAGAAAUUCUACCAGUGU | Ordered from IDT | cleavage, EMSA | Figure 7,8 |
| 6 | target RNA | GUUACCACAGAAAUUCUACCAGUGU | Ordered from IDT | HDX | Figure 2,5,6,7 |

Table S2. HDX-MS parameters

| **Data Set** | ***Di*Cas7-11** | ***Di*Cas7-11 +crRNA(37)** | ***Di*Cas7-11 +crRNA(57)** | ***dDi*Cas7-11 +crRNA(37)** | ***dDi*Cas7-11 +crRNA(37)+ tgRNA** |
|---|---|---|---|---|---|
| HDX reaction details | 20 mM HEPES-NaOH, 150 mM NaCl, 1 mM DTT, pH 7.5 @ 22 °C | | | | |
| HDX time course (min) | 0.5,1,2 | | | | |
| HDX control samples | We use the highest exchanging peptides as maximally deuterated controls | | | | |
| Back-exchange (mean / IQR) | 21.2%/8% | 21.2%/8% | 21.2%/8% | 17.2%/9.3% | 17.2%/9.3% |
| # of Peptides | 194 | 194 | 194 | 212 | 210 |
| Sequence coverage | 94.3% | 94.3% | 94.3% | 93.7% | 90.8% |
| Average peptide length / Redundancy | 14.7/1.78 | 14.7/1.78 | 14.7/1.78 | 14.9/1.97 | 14.7/1.92 |
| Replicates (biological or technical) | 3 technical replicates each | | | | |
| Repeatability (average standard deviation) | 0.068 | 0.076 | 0.083 | 0.095 | 0.087 |
| Significant differences in HDX (delta HDX > X D) | 0.5 D (99%CI) | | | | |

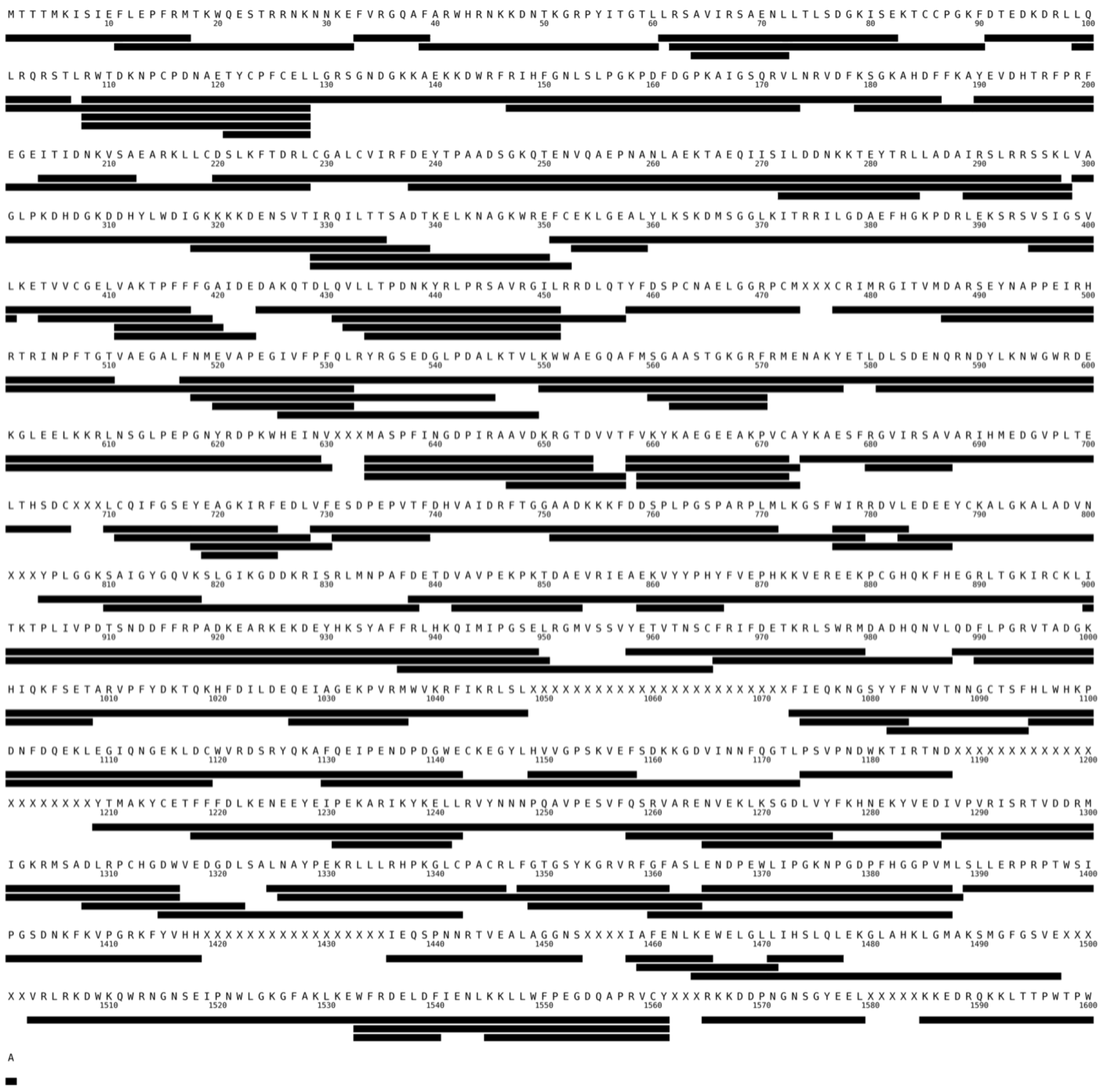

**Supplementary Figure S1.** Coverage map showing all peptides detected and analyzed across all datasets of WT *Di*Cas7-11 from apo as well as $crRNA_{37}$ and $crRNA_{57}$ addition experiments. X's indicate amino acids that were not observed in any state.

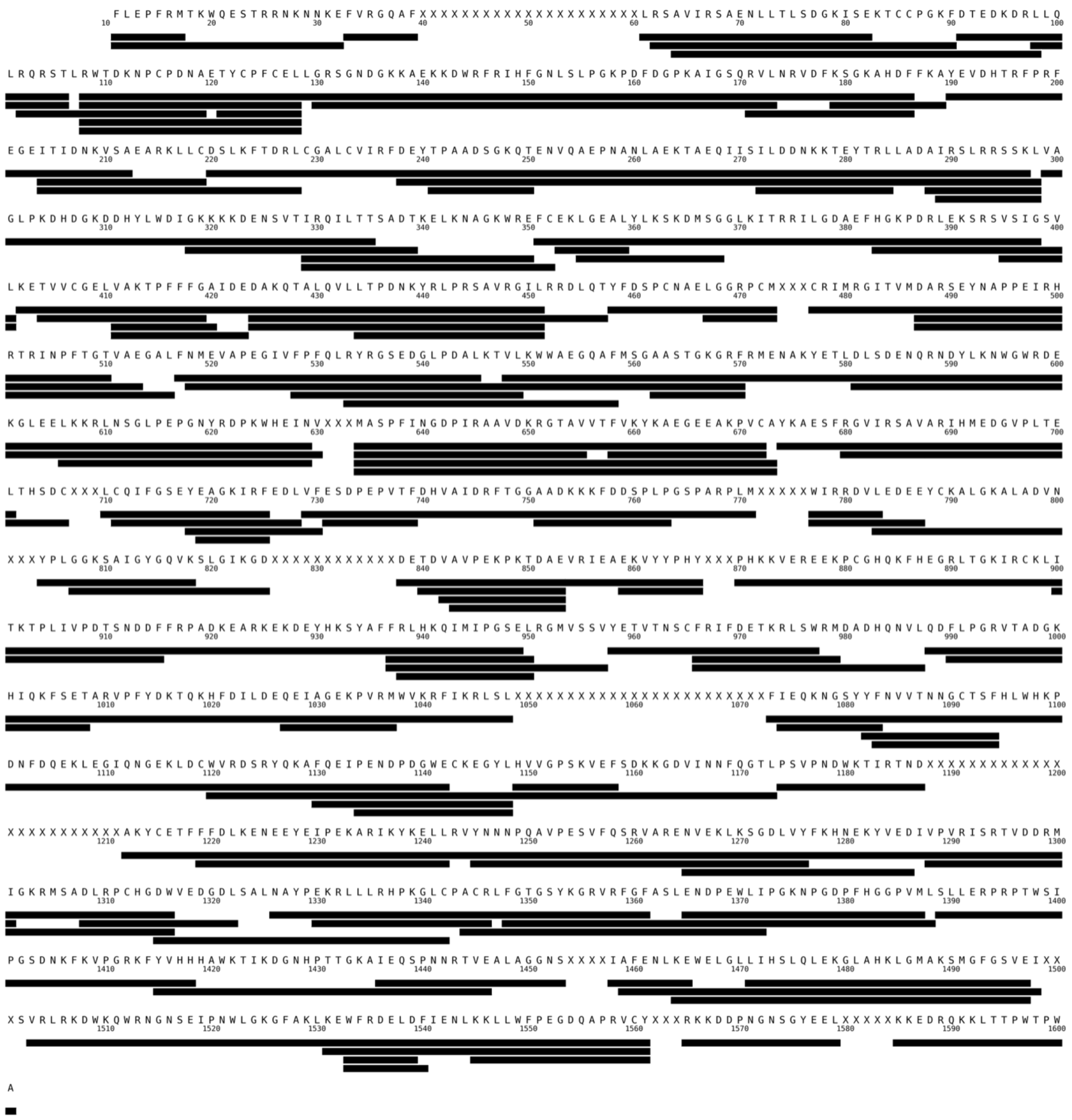

**Supplementary Figure S2.** Coverage map showing all peptides detected and analyzed across datasets of d*Di*Cas7-11-crRNA and d*Di*Cas7-11-crRNA-tgRNA experiments. X's indicate amino acids that were not observed in any state.

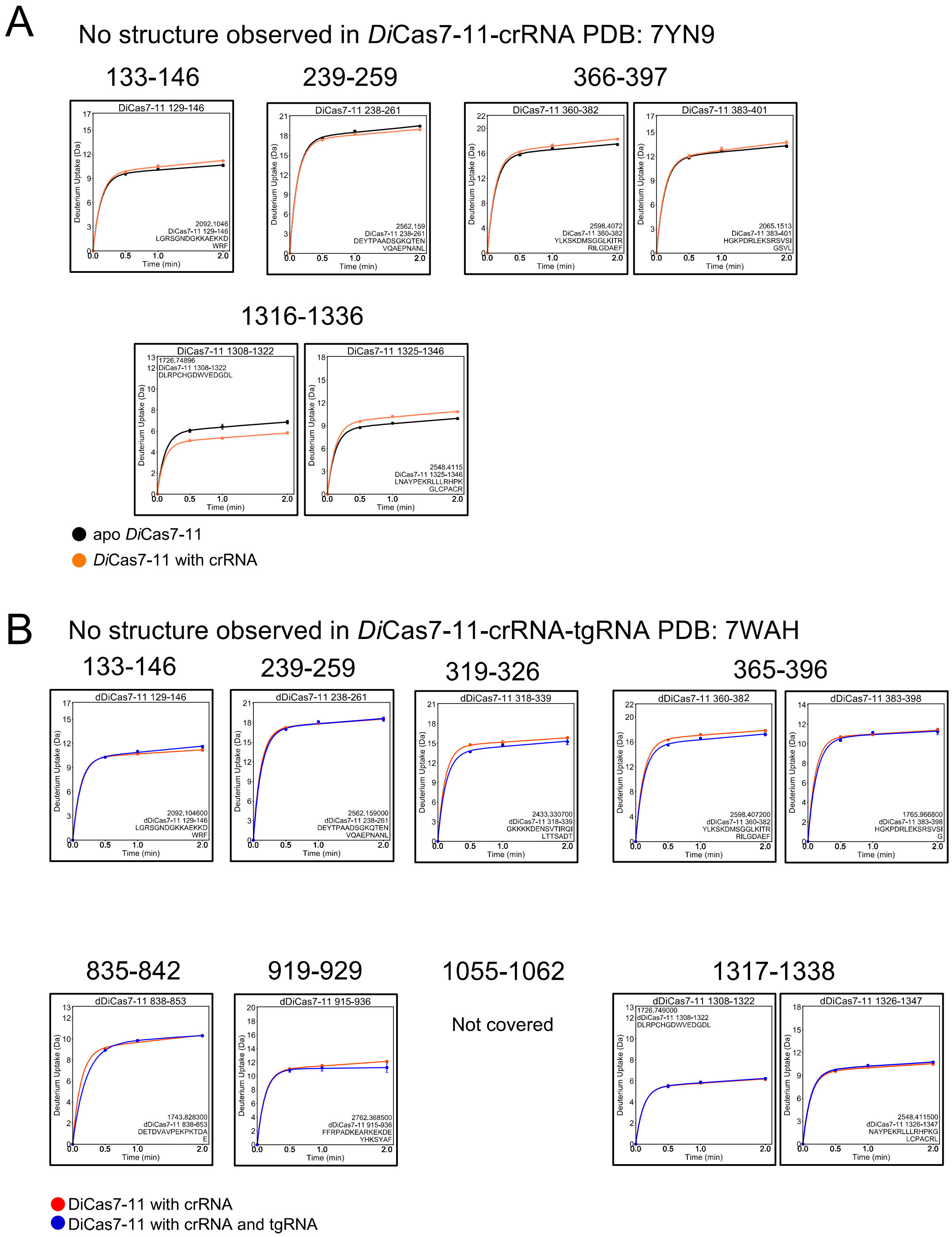


**Supplementary Figure S3.** Uptake plots of regions not covered in structures of (**A**) *Di*Cas7-11-crRNA (PDB:7YN9) (**B**) *Di*Cas7-11-crRNA-tgRNA (PDB:7WAH)

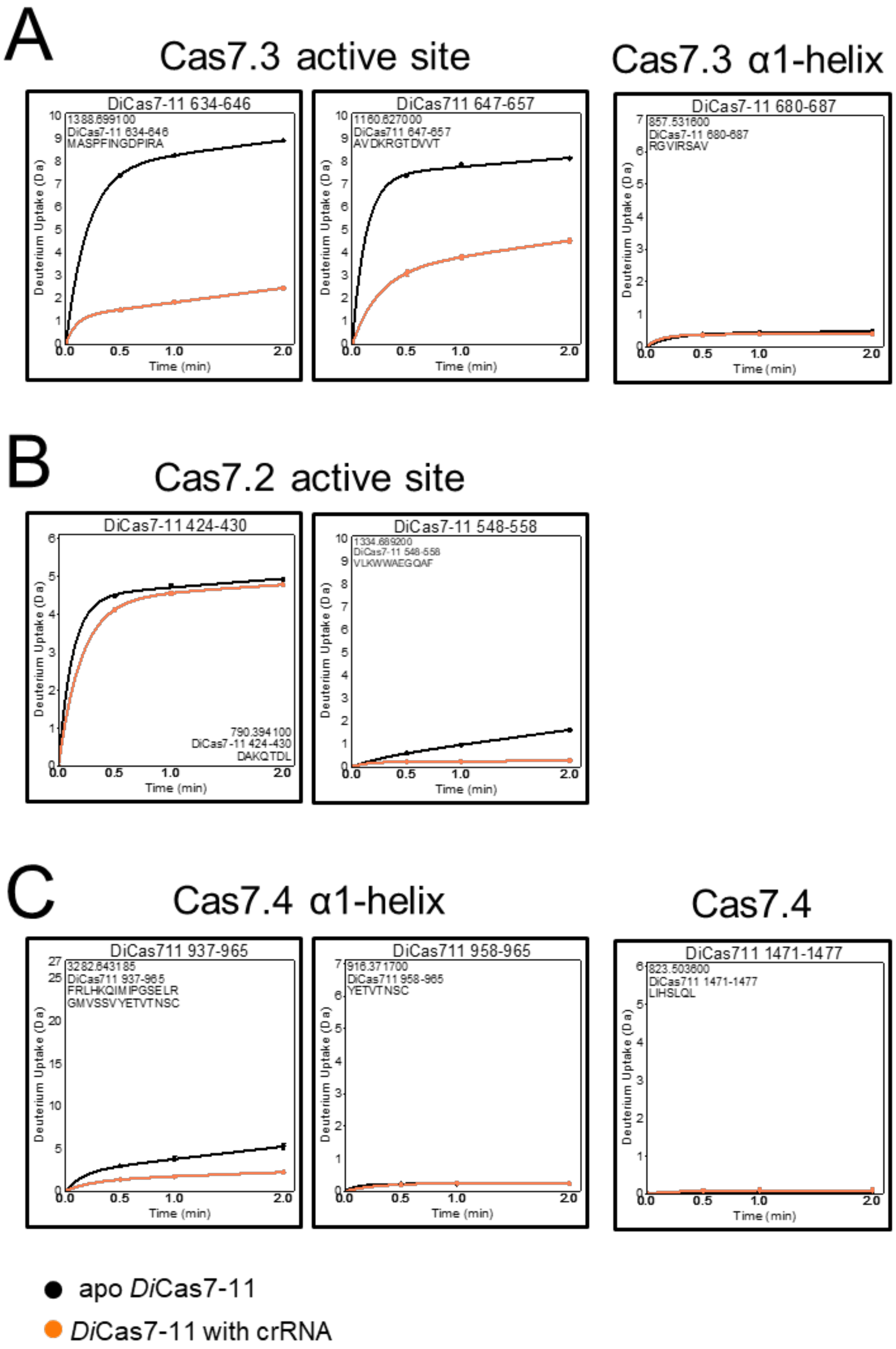


**Supplementary Figure S4.** Additional HDX-MS uptake plots pertaining to **Figure 4** corresponding to (**A**) Cas7.3 active site and α1-helix, (**B**) Cas7.2 active site, (**C**) Cas7.4 α1-helix, and a disordered loop of Cas7.4 that interacts with a nearby flipped base.

### Cas11 domain

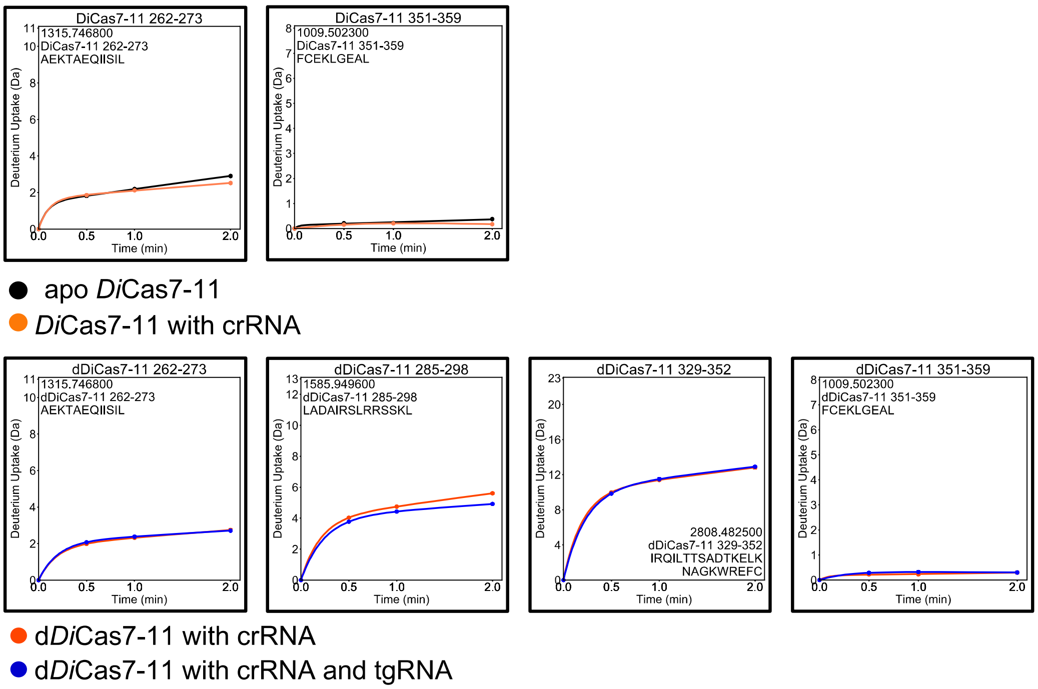


**Supplementary Figure S5.** Additional HDX-MS uptake plots covering the Cas11 domain pertaining to **Figure 5**. Together with main figure plots, the entire Cas11 domain is covered. These regions showed no change from crRNA or tgRNA binding.

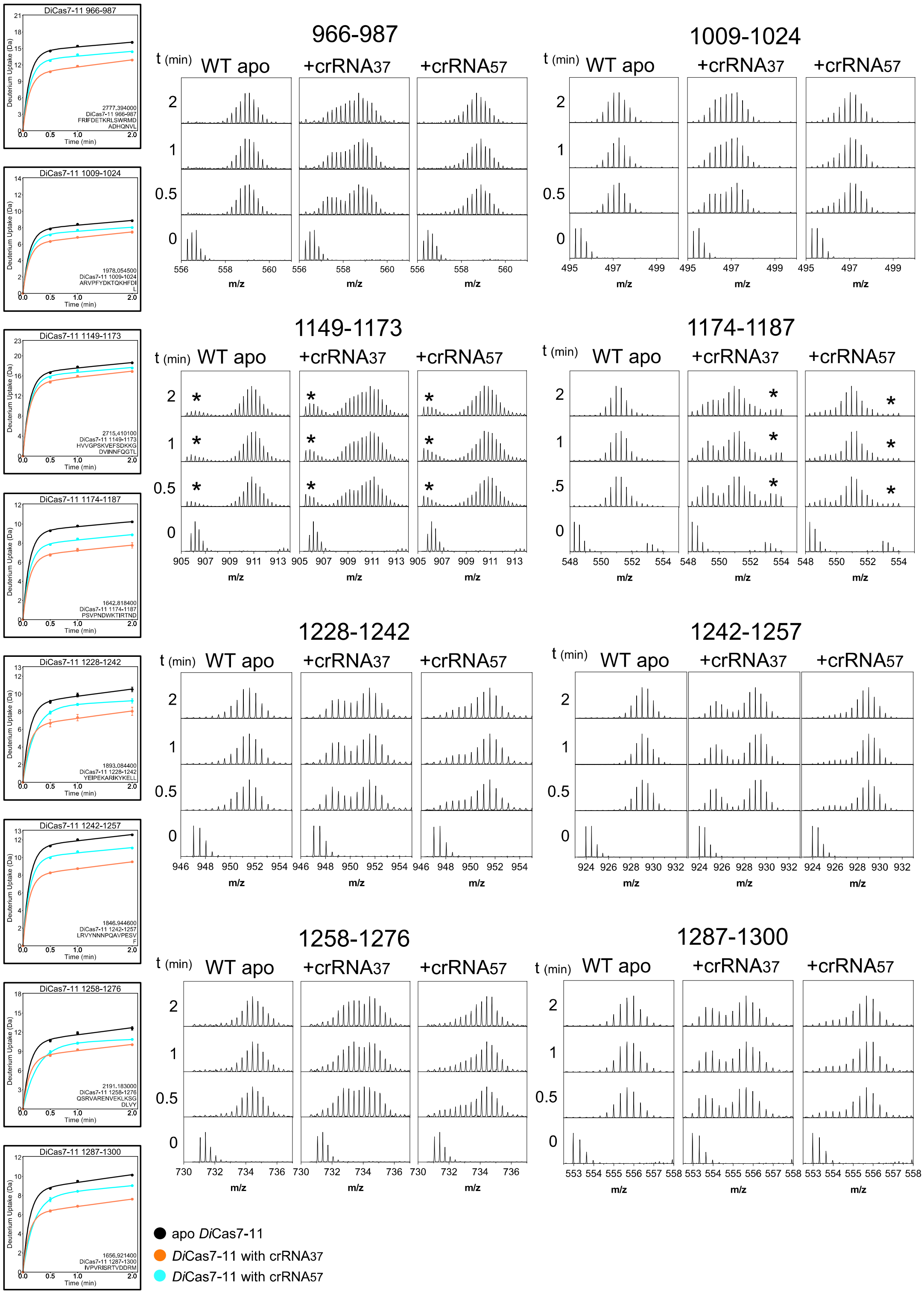


**Supplementary Figure S6.** Raw mass spectra of the INS domain showing bimodal envelope distributions upon crRNA binding. $crRNA_{57}$ showed a bimodal envelope distribution indicative of a more open INS domain. These spectra refer to regions colored green in **Figure 7F**. Mass envelopes marked by asterisks do not belong to the peptide and were not included in data analysis.

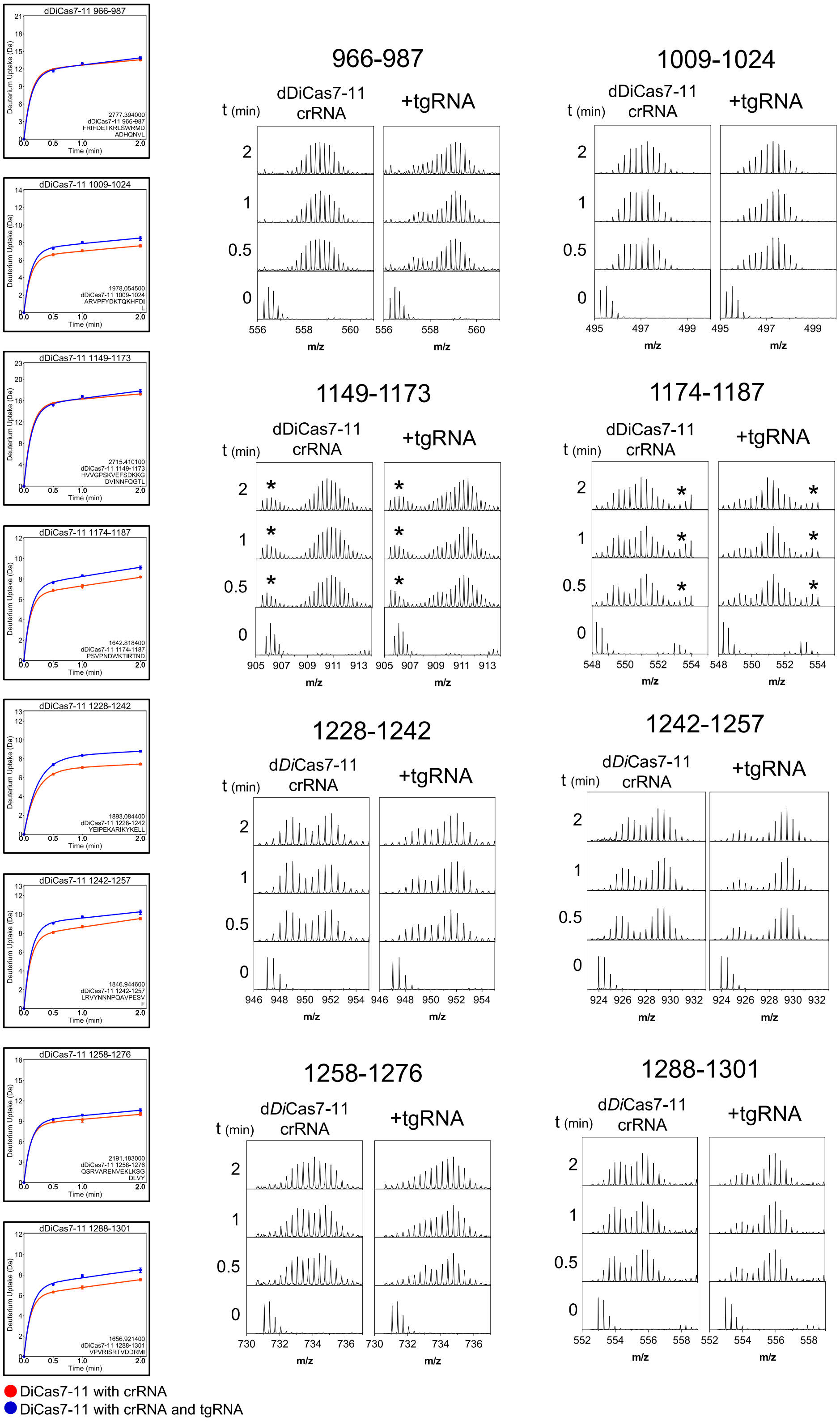


**Supplementary Figure S7.** Raw mass spectra of the INS domain showing a change in bimodal envelope distributions upon tgRNA binding, suggesting a more open INS domain. These spectra refer to regions colored green in **Figure 7I**. Mass envelopes marked by asterisks do not belong to the peptide and were not included in data analysis.

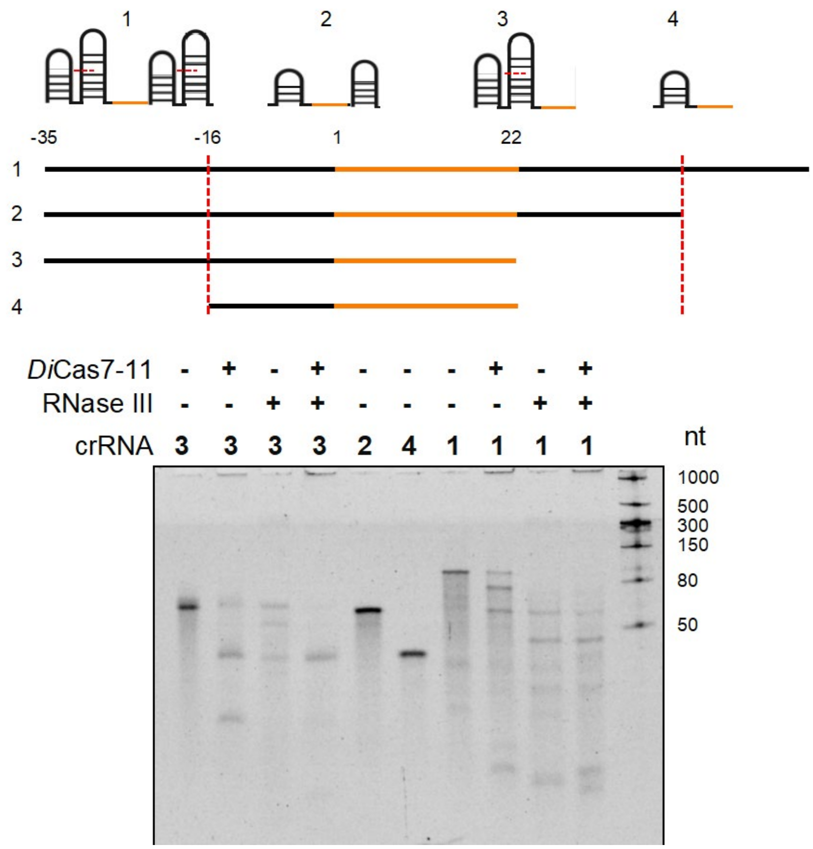


**Supplementary Figure S8.** Processing of pre-crRNA's with *Di*Cas7-11 and RNase III.

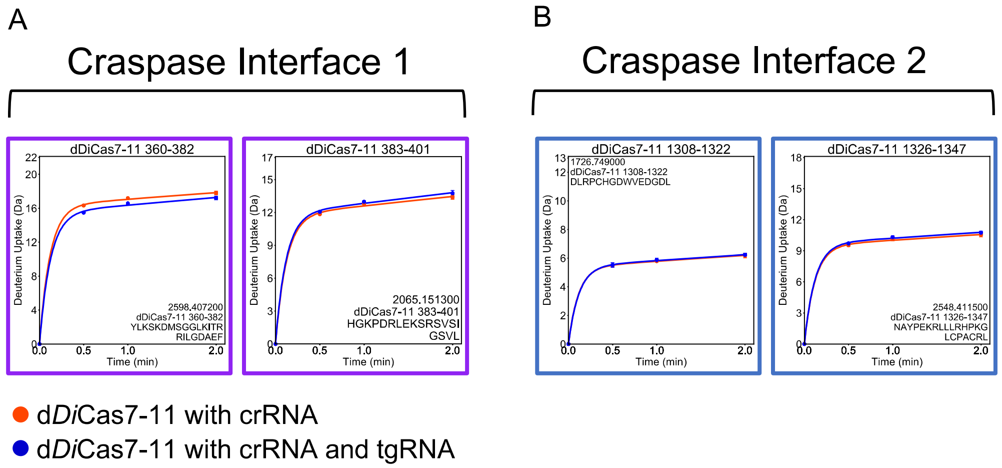


**Supplementary Figure S9.** No changes observed in the Craspase interfaces of d*Di*Cas7-11-crRNA upon tgRNA binding. (A) Craspase interface 1 in d*Di*Cas7-11-crRNA does not change upon tgRNA binding. This interface remains highly solvent accessible under all conditions. (B) Craspase interface 2 in d*Di*Cas7-11-crRNA does not change upon tgRNA binding.
